# Hypoxia inducible factors regulate Pneumovirus replication by enhancing innate immune sensing

**DOI:** 10.1101/2025.08.12.669926

**Authors:** Jiyeon Ha, Parul Sharma, Sammi Ta, Senko Tsukuda, James M. Harris, Rebekah Penrice-Randal, Eleanor Bentley, Adam Kirby, Daniele F. Mega, David A. Matthews, Peter Balfe, Jan Rehwinkel, Anja Kipar, James P. Stewart, Jane A. McKeating, Peter A.C. Wing

## Abstract

The immune mechanisms responsible for protection and pathogenesis in pneumoviral infection are not well defined. We demonstrated that pharmacological activation of the hypoxic inducible factor (HIF) signalling axis using Daprodustat limited viral replication through enhanced immune signalling. Transcriptomic analysis revealed HIF augmented activation of innate immune response genes, including interferon-stimulated gene 15 (*Isg15*), in the lung and spleen of mice infected with pneumonia virus of mice (PVM). In human respiratory syncytial virus (hRSV) infected airway epithelial cells, Daprodustat inhibited viral replication and enhanced ISG15 expression in a HIF-dependent manner. Importantly, inhibition of type I interferon signalling or the RIG-I sensing pathway abrogated the antiviral activity of HIF. Moreover, Daprodustat increased interferon signalling in response to viral RNA, suggesting that HIF inhibits pneumovirus replication through enhancing viral RNA sensing. Mechanistically, Daprodustat reduced N6-methyladenosine modification of viral RNA through upregulation of RNA demethylases, promoting detection by innate immune sensors. This study highlights the intricate interplay between hypoxia and antiviral immunity and offers valuable insights into pneumovirus-host interactions and potential therapeutic interventions.

**SIGNIFICANCE STATEMENT:** This study explores the role of the hypoxic inducible factor (HIF) signalling pathway in limiting pneumoviral infections, such as respiratory syncytial virus (hRSV) and pneumonia virus of mice (PVM). Using Daprodustat, a clinically approved activator of HIF signalling we demonstrated that stimulation of this pathway enhanced innate immune signalling and limited viral replication. Transcriptomic analysis showed that HIF promoted innate immune response genes in mice infected with PVM and human airway epithelial cells infected with hRSV. Importantly, inhibition of type I interferon signalling or the RIG-I sensing pathway abrogated the antiviral activity of HIF. We uncovered a role for HIF to regulate the levels of N6-methyladenosine modification of viral RNA transcripts, resulting in the increased activation of nucleic acid sensing.

## INTRODUCTION

Pneumoviruses such as human respiratory syncytial virus (hRSV) cause lower respiratory tract illness in children(1), immunocompromised adults and the elderly(2). hRSV-induced bronchiolitis and pneumonia in infancy can lead to long-term respiratory sequelae, such as asthma(3). Most infections are mild, however some infants develop a life-threatening inflammatory disease of the lower airways, the underlying causes of which are unknown. In low-middle-income countries hRSV is the second most common cause of infant death after malaria(4, 5). Treatment options are limited, although the recent approval of hRSV-fusion protein vaccines for the elderly and pregnant women(4) provides opportunities to limit disease burden. Like other viruses, hRSV evolves resistance to immune and antiviral therapies(6–8), highlighting the need for a range of therapeutic approaches.

The airway epithelium is the first point of contact for hRSV, where infection is sensed by pattern recognition receptors (PRR), including Toll-like receptors (TLRs) and retinoic acid-inducible gene 1 (RIG-I)-like receptors (RLRs), which act in concert to trigger type-I interferon (IFN) expression to limit viral replication and spread(9). However, hRSV has developed effective countermeasures to circumvent host innate sensing and encodes several proteins that perturb multiple aspects of immune signalling. The effectiveness of these strategies is highlighted by the low to undetectable levels of type I IFN in nasal washes collected from infants with severe hRSV infection, in stark contrast to influenza A or parainfluenza virus infection(10–12). Mononuclear cells obtained from infants with severe bronchiolitis express low levels of IFN-α(12) and *ex vivo* infection of human macrophages or peripheral blood mononuclear cells with hRSV leads to modest type I IFN expression(13). Somewhat paradoxically, hRSV infected lung epithelial cells from infants have been reported to express high levels of IFN-stimulated genes such as ISG15, a ubiquitin-like protein with immunomodulatory properties, mediated through a process known as ISGylation(14). Notably, an association between high levels of hRSV RNA and ISG15 expression in nasal aspirates was reported in infected infants, highlighting the potential use of ISG15 as a biomarker of hRSV-induced inflammation(15).

Among the various hRSV proteins involved in immune evasion, the best characterised are the non-structural (NS) 1 and 2 proteins(16). The NS1-2 open reading frames (ORFs) are positioned proximal to the 3’ leader region within the viral genome, making them the earliest expressed and most abundant viral transcripts within an infected cell(17, 18). Indeed, these two NS proteins distinguishes hRSV and PVM from other members of the *Mononegavirales*. NS1 and NS2 antagonism of type I and III IFN production has been extensively studied and is supported by the attenuation of viral replication in strains that lack either isoform in both *in vivo* and *in vitro* replication models(19–21). Notably these proteins disrupt the interaction between RIG-I and mitochondrial antiviral signalling protein (MAVS), either by direct binding to MAVS(22) or by targeting TRIM25(23). The hRSV nucleoprotein (N), an integral component of the viral nucleocapsid, forms decameric rings around the single stranded (ss) RNA genome(24, 25) and limits the recognition by cellular PRRs. N also sequesters innate signalling proteins into multimeric biomolecular condensates termed inclusion bodies (IBs) that are formed by the N and phosphoprotein (P) that drives liquid phase separation to generate IBs(26–28). Notably, these cytoplasmic structures are sites of viral replication, where viral proteins and replicase complexes are located(27). Recent work has shown that IB-associated granules concentrate nascent viral RNA complexes and components of the eukaryotic translation machinery to drive translation initiation(29). In early replication, MAVS and MDA5 are sequestered to small IBs through interactions with N, thereby inhibiting their antiviral function(s)(30), demonstrating how hRSV perturbs the spatial distribution of host antiviral defences. The multifaceted approach by which hRSV evades and manipulates the host immune responses highlights the complexity of virus-host interactions and underscores the challenges in developing effective interventions against this pathogen.

While hRSV employs sophisticated mechanisms to evade host immune responses, our understanding of the cellular factors that influence viral replication and inflammation is more limited. In this context, hypoxia-inducible factors (HIFs) have emerged as important regulators of both viral infection and inflammatory responses in the lower respiratory tract(31, 32). HIFs are conserved transcription factors that define the response of the transcriptome and metabolome to changing oxygen levels(33). The three isoforms (HIF-1α, HIF-2α and the lesser studied HIF-3α) are regulated by prolyl-hydroxylase domain (PHD) enzymes whose activity depends on oxygen, iron and 2-oxoglutarate. When oxygen is abundant, hydroxylation of two specific prolyl residues in the HIF-α sub-units promote interaction with the von Hippel-Lindau ubiquitin (VHL) E3 ligase leading to ubiquitylation and proteasomal degradation(34–36). When oxygen is limited, this activity is suppressed and HIF-α sub-units dimerise with HIF-1β, forming an active transcriptome complex. Through interactions with cellular importins, these complexes translocate to the nucleus and drive transcription through interaction with a consensus RCGTG(C) motif, also known as hypoxic responsive element (HRE), in the promoter and enhancer regions of responsive genes(37). Additionally, non-hypoxic stimuli, including metabolic and inflammatory signals, can regulate HIF expression(38, 39). HIF-regulated genes vary between cell types and silencing HIF-1α/-2α reveals their unique transcriptional signatures, highlighting tissue-and isoform-specific responses to these diverse physiological signals(40). Clinically, several PHD inhibitors (PHDi) are licensed for treatment of anemia(41). Our recent studies show a dual role for HIFs in restricting hRSV and SARS-CoV-2 infection and inflammatory responses(42–44), providing opportunities to exploit current licensed HIF-mimetic drugs for the treatment of respiratory infections.

Animal models of human hRSV (hRSV) are limited by the semi-permissive nature of many rodent and non-human primates(45). Chimpanzees are the exception and replicate comparable levels of hRSV to human infants(46, 47), which have been used to evaluate the virulence and protective efficacy of live attenuated vaccine candidates(48). BALB/c mice show intermediate susceptibility to experimental hRSV infection(49), and as such high infectious doses are required to induce clinical signs of disease, such as weight loss, ruffled fur, and hunching (50, 51). One model to study pneumovirus infection is the natural mouse pathogen, pneumonia virus of mice (PVM)(52), which is associated with severe morbidity, weight loss, laboured breathing and mortality(53, 54). The virus replicates predominantly in alveolar and bronchial epithelial cells, inducing increased eosinophil counts in bronchial alveolar lavage samples, followed by a dominant neutrophil response(55, 56). Natural *in vivo* models of viral disease such as PVM provide valuable insights into pneumovirus pathogenesis and serve as a platform to evaluate potential therapeutic interventions.

In this study we explored the impact of HIF signalling on pneumoviral disease by treating PVM-infected mice with the licensed PHDi, Daprodustat to activate HIF signalling. Daprodustat inhibited PVM replication and immune infiltration and promoted pulmonary innate-immune signalling. Using *in vitro* model systems of hRSV infection, we confirmed these findings and demonstrated that the antiviral effect of HIF on hRSV replication is dependent on RIG-I signalling. Collectively these findings highlight how targeting HIF-regulated pathways may offer an alternative approach for treating hRSV infection and the associated immune response.

## RESULTS

### PVM causes a widely disseminated pulmonary infection and innate immune response

To study PVM infection in the lower respiratory tract, female BALB/c mice were inoculated intranasally with PVM the strain J3666 at a dose of 1 x 10^4^ PFU per animal. We selected BALB/c given previous reports that this mouse strain showed increased susceptibility to PVM compared to C57BL/6 mice(52). The infected mice maintained their body weight before showing a rapid loss at 5 days post-infection (dpi) (**Fig.1A**). Infectious virus was detected in the lung tissue at 3 dpi (ca. 5 x 10^5^ PFU/ml) and this declined to ca 1.5 x 10^4^ PFU/ml at 5 dpi (**Fig.1B**). Histological examination and staining for PVM-G protein revealed widespread antigen expression in the lung at 3 dpi, represented by multifocal patches of alveoli with type I and II pneumocytes and respiratory epithelial cells expressing the viral antigen along the luminal border in some bronchioles (**Supp Fig.1A**). The associated histological changes were limited to focal areas of increased interstitial cellularity, with activated type II pneumocytes and mild perivascular mononuclear infiltration (**Supp Fig.1A**). By day 5, viral antigen expression was apparent in the alveoli with large focal areas containing desquamated alveolar macrophages/type II pneumocytes and some leukocytes, consistent with acute pneumonia. There was evidence of leukocyte recruitment into the parenchyma, with leukocyte rolling and mild perivascular accumulation (**Supp Fig.1B**). Minimal pathological changes were observed in either the parenchyma or alveoli of mock infected mice (**Supp Fig.1C**). To determine whether PVM had spread beyond the lungs, we examined the heart, thymus, spleen, liver and kidney, and found negligible evidence for any histological changes or viral antigen expression, in line a with previous report(57).

**Figure 1:**
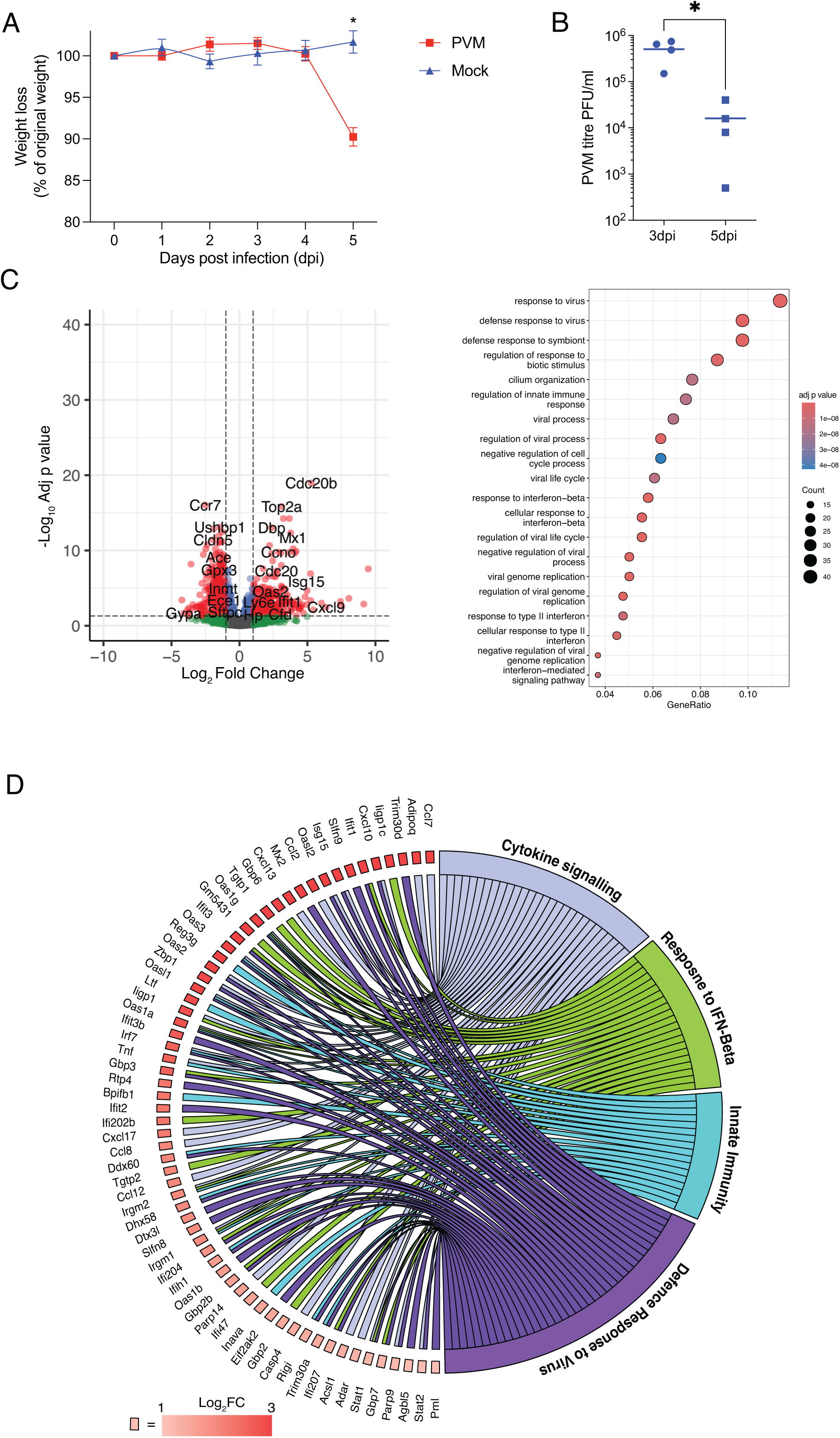
Characterising PVM infection of the lung in mice. **(A)** Mice (n=4) were infected intranasally with 1×10^4^ PFU of PVM-J3666 or PBS and body weight assessed daily up to 5 dpi. Data is mean ± S.D. of n=4 animals. **(B)** Viral titres from lung tissue of infected mice measured at 3 dpi. **(C)** RNA sequencing analysis of uninfected or PVM infected lungs at 3 dpi. The volcano plot depicts differentially expressed genes (DEGs) with thresholds defined as a log_2_ fold change of > 1 or <-1 with an adjusted p value of <0.05 (dashed lines). The symbol colour is derived from-log_10_ of the adjusted p value, where red indicates a log_2_ fold change of > 1 or <-1 with an adjusted p value of <0.05. Up-regulated genes were subjected to gene-ontology analysis, with the top 20 enriched pathways depicted. Size of symbol indicates the number of genes from each pathway enriched in PVM infection. The gene ratio is the relative number of genes in a GO term that are enriched out of the total number of DEGs in the data set. **(D)** Modified circos plot showing the relationship between the up-regulated genes from (C) and enriched GO pathways. Boxes adjacent to the gene labels are coloured according to the log_2_ fold change of the individual gene between the infected and uninfected mice.

To elucidate the underlying molecular mechanisms driving the host antiviral response we sequenced RNA from the lungs of mock or PVM infected animals at 3 dpi before the onset of infection-associated weight loss and distinct pathological changes. PVM induced a substantial change in the pulmonary transcriptome, with gene ontology (GO) analysis showing an enrichment of innate and immune-response pathways, indicative of a widely disseminated pulmonary viral infection in line with previous studies(58) **(Fig.1C)**. We examined the relationship between the differentially expressed genes (DEGs) in response to PVM infection and the enriched immune pathways identified in **Fig.1C**. This highlighted a key set of chemokine (*Cxcl9, Cxcl10, Cxcl13*) and innate immune response genes (*Mx2, Oas2, Tnf, Ifit1 and Isg15*) that were up regulated in PVM infection **(Fig.1D)**, underscoring the robust induction of host immune pathways elicited by PVM infection in the lungs.

### Daprodustat limits PVM infection and associated pathological changes

As HIF signalling was previously shown to modulate the immune response(59) we investigated the therapeutic effect(s) of Daprodustat, a HIF-PHDi, on PVM infection. PVM infected mice were treated 24h post-infection with 30mg/Kg of Daprodustat or vehicle by oral gavage for 3 or 5dpi. Given the decline in infectious viral titre noted between 3 and 5dpi (**Fig.1B)** we examined the impact of Daprodustat on viral replication at 3dpi. The cell types expressing viral antigen were comparable in both groups, however, their frequency was lower following Daprodustat treatment, with fewer bronchioles staining for PVM-G and smaller patches of infected alveolar epithelial cells (**Supp Fig.2A**). This observation was confirmed by a quantitative analysis of PVM-G expression in the lung (**Fig.2A**). To confirm drug activity, we quantified the HIF-target gene endothelin 1 (*Edn1*)(60) in the lung and noted a significantly higher expression in Daprodustat treated animals (**Fig.2A**). To further investigate the effects of Daprodustat, we conducted a transcriptomic analysis of lung tissue from mice at 3dpi. This revealed an enrichment of wound healing, angiogenic and haemostatic pathways (**Fig.2B**), in agreement with our previous work using the HIF-PHDi Roxadustat in a rodent model of SARS-CoV-2 infection(43).

**Figure 2:**
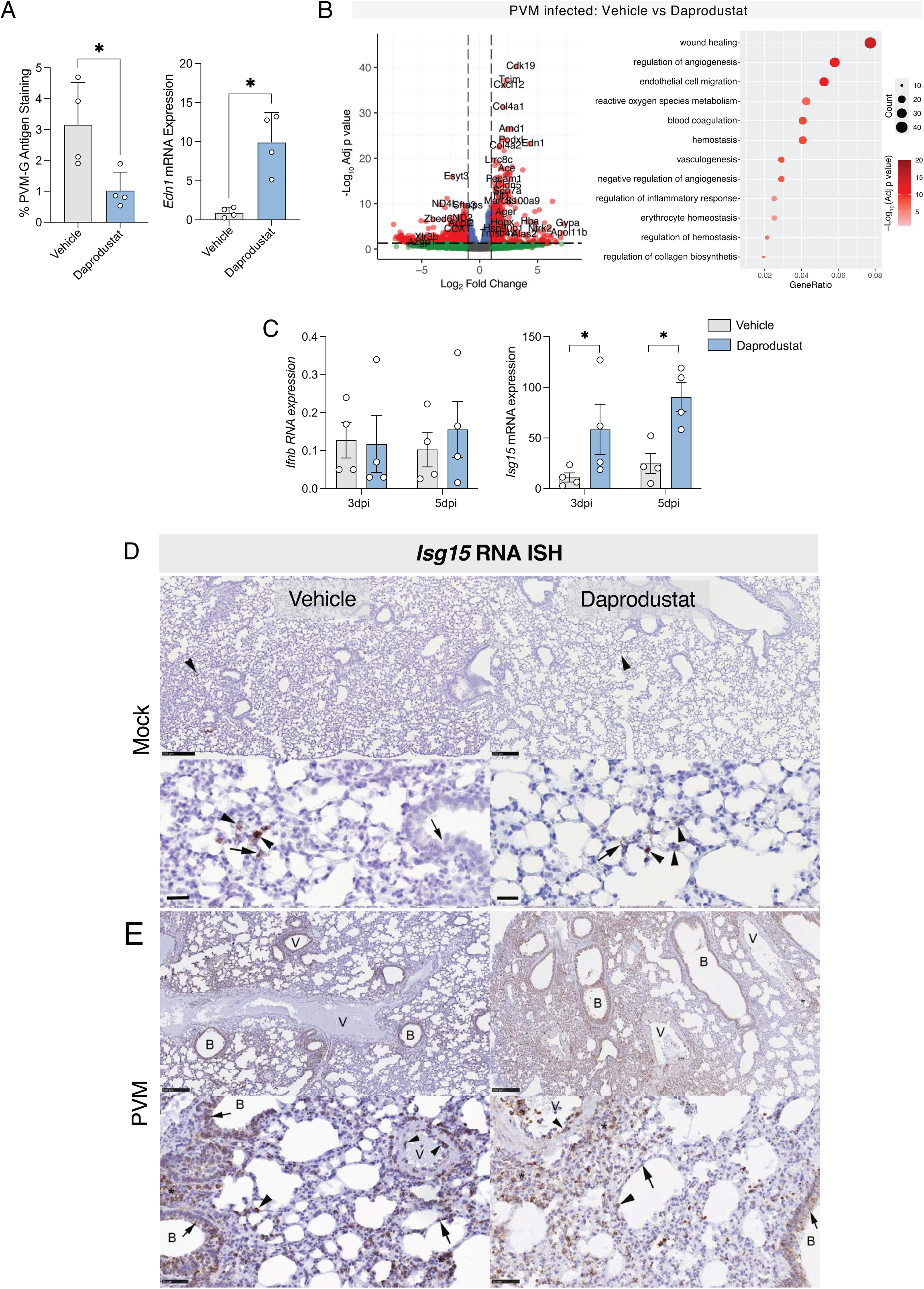
Impact of Daprodustat treatment on PVM infection. **(A)** Quantification of PVM-G expression and RT-qPCR quantification of Endothelin-1 gene expression in lungs from PVM infected mice treated with vehicle or Daprodustat (30mg/Kg) at 3dpi. **(B)** Differential gene expression analysis of Daprodustat-treated, PVM-infected animals compared to infected mice treated with vehicle. DEGs are defined as a log_2_ fold change of > 1 or <-1, with an adjusted p value of <0.05. Up-regulated genes were used for gene-ontology analysis, with the top 20 enriched pathways depicted. **(C)** Quantification of *Ifnb* and *Isg15* mRNA by RT-qPCR from lung tissue harvested at 3 and 5dpi from vehicle and drug treated groups. Data is expressed relative to the *Actb* housekeeper gene. **(D)** Detection of *Isg15* mRNA expressing cells in the lungs of vehicle (left column) and Daprodustat treated (30 mg/Kg, right column) mock infected mice. The rare small patches of positivity in type I (arrows) and type II (arrowheads) pneumocytes in both treatment groups are annotated. A low signal was detected in some respiratory epithelial cells in the bronchiole (small arrow). **(E)** *Isg15* mRNA expression in PVM infected mice treated with vehicle or Daprodustat at 5dpi. Widespread *Isg15* mRNA expression is seen after PVM infection in the bronchioles and parenchyma. Small arrows highlight expression in bronchiolar epithelial cells with large arrows and arrowheads denoting strong expression in type I and II pneumocytes, respectively. B: bronchiole; V: vessel. Scale bars represent 50µm.

Treatment of infected mice with Daprodustat resulted in a significant increase in regulatory inflammatory gene expression such as *Cxcl12* and *Tgfb1* (**Fig.2B**), prompting further investigation of the composition and extent of immune cell infiltration in the lung tissue. We selected samples at 5dpi for detailed histological analysis, considering the more substantial pathological changes observed at this timepoint (**Supp Fig.1B**). Examination of these tissues revealed focal areas of consolidation in the lung, with desquamation of alveolar epithelia/alveolar macrophages and leukocyte infiltration in the alveolar lumina (**Supp Fig.2B**). Vehicle-treated animals exhibited emigration and perivascular accumulation of mononuclear leukocytes around blood vessels, in contrast to the Daprodustat treated mice (**Supp Fig.2B**). Given the increased expression of chemokine and cytokine genes in response to PVM infection from our earlier transcriptomic analysis **(Fig.1C-D),** we assessed the extent of leukocyte recruitment and composition of perivascular infiltrates in both vehicle and drug treated groups. Lung sections were immunostained to visualise T cells (CD3+), monocytes/macrophages (Iba1+) and neutrophils (Ly6G+). Vehicle treated mice showed extensive monocyte/macrophage recruitment into the lung parenchyma (**Supp Fig.2B**). T cells were less abundant, representing individual cells in the lumen of capillaries and in inflammatory infiltrates (**Supp Fig.2B**). In contrast, in the PVM infected-Daprodustat treated mice, there was limited evidence of monocyte/macrophage recruitment and infiltration, accompanied by a few T cells (**Supp Fig.2B).** Neutrophils were observed to be more prevalent and were identified within capillaries and in regions of alveolar damage and infiltration, with comparable quantities in both groups (**Supp Fig.2B**). These findings highlight the differential immune cell recruitment and infiltration patterns in PVM-infected mice treated with Daprodustat compared to vehicle-treated controls, suggesting an immunomodulatory role for HIFs in the regulation of inflammatory response during viral infection.

To assess the impact of HIF on the host response to PVM, we performed targeted gene expression analysis of *Isg15* and *Ifnb* transcripts by RT-qPCR from vehicle and Daprodustat treated groups. Daprodustat augmented *Isg15* expression but had a minimal effect on *Ifnb* **(Fig.2C)**. We also examined the spatial distribution of *Isg15* transcripts in the tissue samples using RNA *in-situ* hybridisation (ISH), noting limited expression in the lungs of uninfected mice, irrespective of Daprodustat treatment **(Fig.2D).** However, in the spleen, moderate numbers of *Isg15* positive cells were detected in the red pulp of uninfected animals treated with vehicle, with a modest increase following Daprodustat treatment **(Supp Fig.3A)**. Extensive *Isg15* mRNA expression was observed in the lungs of the PVM infected mice at 5 dpi **(Fig.2E)** with alveolar and bronchiolar epithelial cells, as well as infiltrating leukocytes and vascular endothelial cells, showing high levels of *Isg15* expression which appeared more extensive with Daprodustat treatment **(Fig.2E)**. In PVM infected mice the frequency of *Isg15* mRNA expressing cells in the splenic red pulp was substantially increased and this was more pronounced following Daprodustat treatment **(Supp Fig.3B)**. To assess the cell types expressing *Isg15* in the lung, sections were co-stained with the monocyte/macrophage marker Iba1 or the neutrophil marker Ly6G, demonstrating *Isg15* expression in monocytes, infiltrating/alveolar macrophages and neutrophils, respectively **(Supp Fig.3C)**. Together, these tissue-specific and infection-dependent effects of Daprodustat on *Isg15* expression highlight a role for HIFs in regulating the innate immune response to PVM infection.

### Daprodustat promotes innate immune gene expression in PVM infection

Building on our histopathological analysis of the lung tissue, we examined the broader transcriptomic changes induced by Daprodustat during PVM infection. Daprodustat treatment resulted in an enrichment of the innate-response pathways in PVM infected mice, with higher gene-ratios noted for each GO pathway **(Fig.3A)**. We therefore interrogated our RNA sequencing data with a focus on a consolidated list of innate response genes based on the significantly enriched innate immune GO pathways observed in vehicle-treated-PVM infected mice. Hierarchical clustering showed a significant increase in the expression of RNA editing enzymes, *Adar* and *Apobec3*, type I IFN response genes such as *Ifit1, Ifit2, Oas2, Oas3 and Isg15,* and PPR genes, including *Ddx58* (encoding RIG-I) and *Tlr3*, in PVM infected animals compared to mock-infected controls **(Supp Fig.4)**. Notably, Daprodustat treatment enhanced the expression of innate genes, including an array of IFN genes, that were not observed in PVM infection alone, such as *Ifng, Ifna1, Ifna4, Tlr9,* and *Trim38* **(Supp Fig.4A).** The increased expression of toll-like receptors such as *Tlr9* and *Tlr3* in Daprodustat treated infected mice, suggests that HIFs may prime the innate immune response to detect a broad range of viral nucleic acids across the endosomal pathway. Mapping the potential cell types in the lung using digital cell quantification (61) suggests that Daprodustat modulates the abundance of monocytes, dendritic and NK cells **(Supp Fig.4B-C)**. Notably we observed reduced macrophage signatures in line with our immunohistochemical analysis of infected lung tissue **(Supp Fig.2B)**. Collectively these findings suggest a wider impact of HIF regulation of the innate immune system to detect viral pathogens.

**Figure 3:**
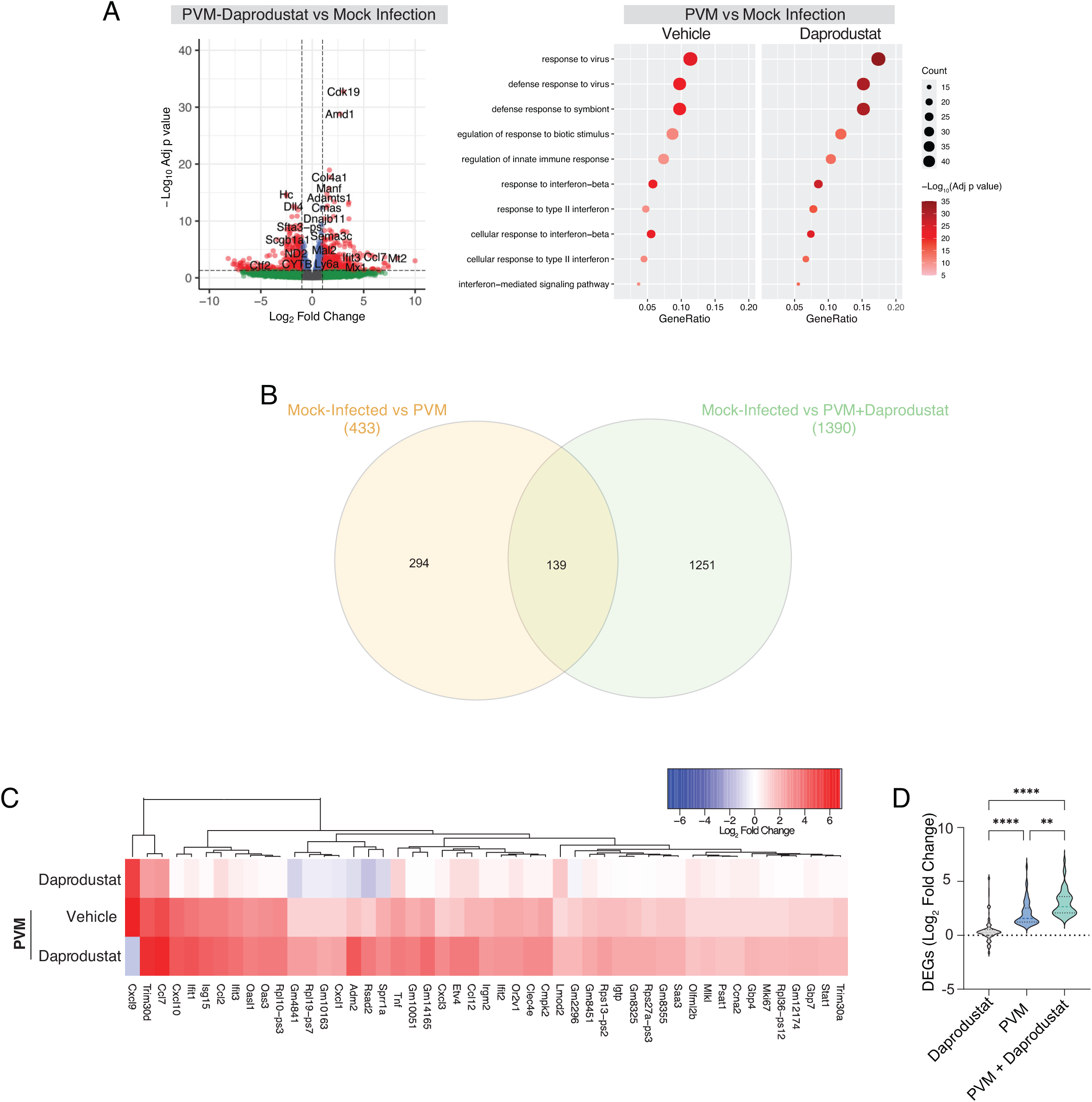
Daprodustat augments innate immune responses to PVM. **(A)** Differential gene expression analysis of Daprodustat-treated PVM-infected animals compared to the vehicle control group. Volcano plot indicates significant DEGs as defined as a log_2_ fold change of > 1 or <-1, with an adjusted p value of <0.05. Comparison of enriched GO pathways in uninfected and PVM-infected animals treated with vehicle or Daprodustat. Size of symbol indicates the number of genes from each pathway enriched by vehicle or Daprodustat treatment. The symbol colour is derived from-log_10_ of the adjusted p value. The gene ratio is the relative number of genes in a GO term that are enriched out of the total number of DEGs for each dataset. **(B)** Venn diagram showing the intersection between up-regulated DEGs in PVM and PVM-Daprodustat treated animals compared with uninfected animals. **(C)** Hierarchical clustering of the 58 common innate immune genes (derived from B) in Daprodustat treated uninfected mice and PVM infected mice treated with vpehicle or Daprodustat. Colours are derived from the log_2_ fold change in expression relative to uninfected, untreated animals, where red indicates a log_2_ fold change of > 1 or <-1 with an adjusted p value of <0.05. **(D)** The log_2_ fold changes from the entire data set from C were compared by ANOVA; p <0.01 = ** and p<0.0001 = ****.

Analysis of the lung transcriptomic data from PVM-infected animals treated with vehicle or Daprodustat, identified a group of 139 genes with significantly increased expression relative to mock-infected animals (**Fig.3B).** To focus our analysis on innate response genes we narrowed our investigation to a list of 58 genes, previously identified as involved in the innate immune response. Daprodustat significantly enhanced expression of this gene set in PVM infection and, importantly, we noted negligible modulation of these genes in uninfected mice treated with Daprodustat, indicating that PVM infection was a prerequisite to observe these changes in innate gene expression (**Fig.3C-D)**.

### Daprodustat inhibits hRSV infection

We investigated whether the elevated expression of innate antiviral genes such as *Isg15, Ifnl1, Ifna-1* and *-4* in the Daprodustat treated, PVM infected mice extended to hRSV. Air-liquid-interface cultures of human primary bronchial epithelial cells (ALI-PBEC) were used to assess the impact of Daprodustat on innate response gene expression following hRSV infection. Daprodustat treatment resulted in a significant reduction in hRSV-N gene expression and a concomitant increase in the HIF-target gene N-myc downstream-regulated gene 1 (NDRG1) expression **(Fig.4A)**. In line with our PVM infected lung RNA-seq **(Supp Fig.4)**, Daprodustat increased innate antiviral gene expression **(Fig.4)**. To extend this observation we infected Calu-3 lung epithelial cells with increasing doses of hRSV followed by Daprodustat treatment or cultured under low oxygen conditions (1% O_2_) previously reported to induce HIF expression(42). Both treatments significantly reduced hRSV-N gene and protein expression **(Fig.4B)**. Stabilisation of both HIF-1 and HIF-2α isoforms and induction of NDRG1 confirmed Daprodustat activity **(Fig.4C)**. We detected increased levels of ISG15 in hRSV infected cells treated with Daprodustat, confirming our observations with the ALI-PBECs and the *in vivo* data obtained from PVM infected mice **(Fig.4C)**. Comparing the impact of either Daprodustat or hypoxia on PVM replication in baby hamster kidney (BHK) cells, which support PVM replication(62), showed that both treatments significantly reduced PVM N and NS2 gene expression, as well as the infectious viral load **(Supp Fig.5A-C)**. To assess whether hRSV infection induced IFN signalling we incubated the extracellular media from infected HEp-2 cells, a well described hRSV-permissive cell line(63), with HEK-293 cells expressing a luciferase-based interferon-stimulated response element (ISRE) reporter construct. hRSV activated the ISRE reporter, with Daprodustat significantly enhancing this effect, indicative of increased type I IFN production **(Supp Fig.5D)**. To assess whether Daprodustat inhibition of hRSV replication is dependent on innate immune signalling we evaluated the antiviral capacity of Daprodustat in Vero cells, which lack the ability to produce endogenous IFN(64), but exhibit comparable responses to Daprodustat as assessed by NDRG1 expression (**Supp Fig.5E**). Vero cells were highly permissive for hRSV replication, however the inhibitory activity of Daprodustat was diminished compared to Calu-3 cells, suggesting that IFN signalling is a major component of the antiviral activity of Daprodustat **(Fig.4D, Supp Fig.5E)**.

**Figure 4:**
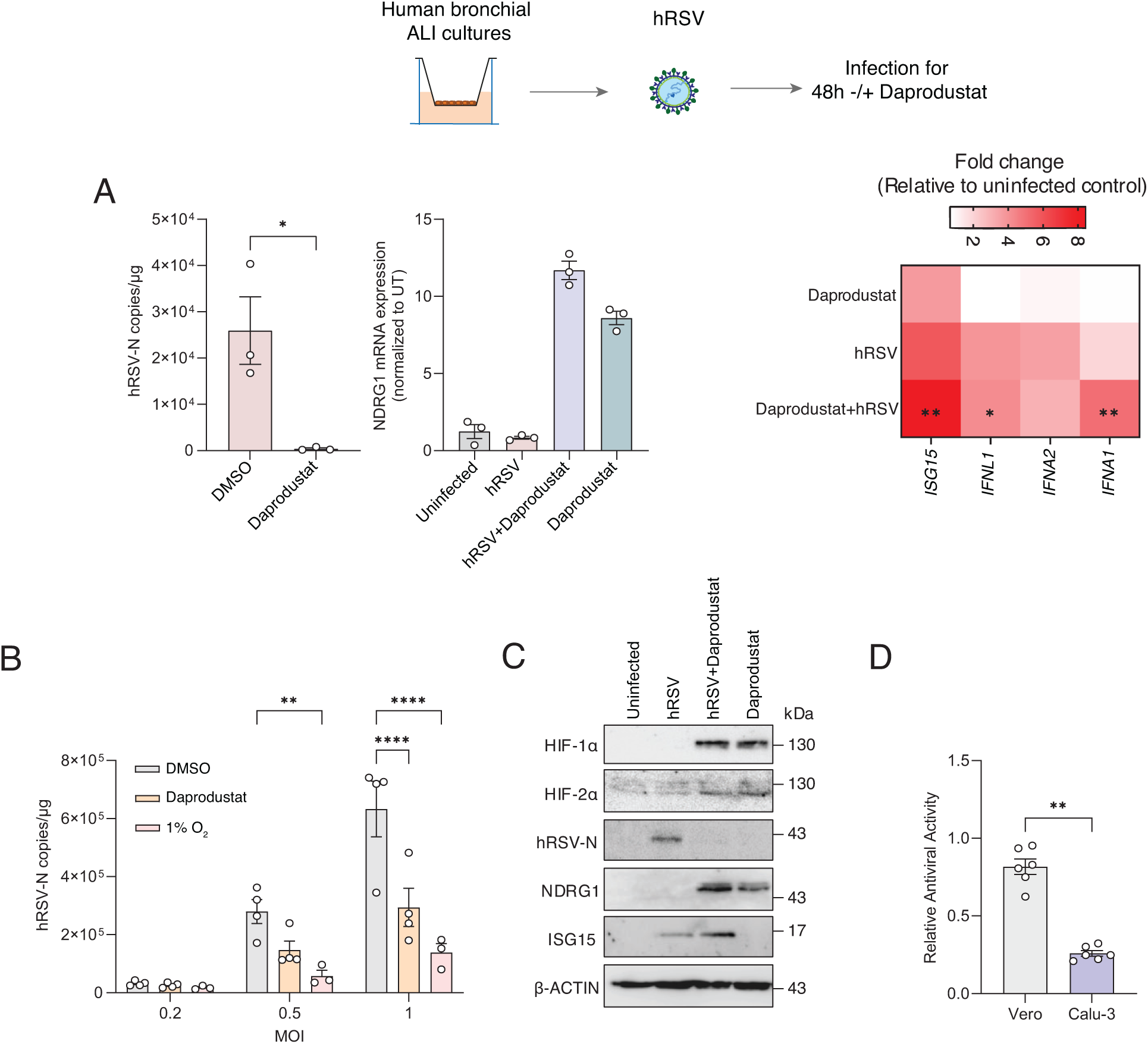
Daprodustat and low oxygen inhibit hRSV infection and increase ISG15 expression. **(A)** ALI-differentiated PBECs from 3 donors were infected with hRSV at an MOI of 1 and treated with Daprodustat (50µM) for 48h. hRSV-N and expression of *NDRG1, ISG15, IFNL1, IFNA1 and IFNA2* were assessed by RT-qPCR. **(B)** Calu-3 were infected with hRSV at a range of MOIs and treated with 50µM of Daprodustat or 1% O_2_ for 48h. hRSV-N was quantified by RT-qPCR with statistical significance assessed by ANOVA. **(C)** Protein expression of HIF-1α and 2α, hRSV-N, NDRG1 and ISG15 in Calu-3 cells treated for 48h with Daprodustat with or without hRSV infection. **(D)** Vero or Calu-3 cells were infected with hRSV (MOI 1) and treated with 50µM of Daprodustat for 48h. Viral transcripts were quantified by RT-qPCR and compared relative to their DMSO control to determine the relative antiviral effect of the drug treatment. Unless otherwise stated, plotted points indicate independent biological replicates and shown as mean ± SD. Statistical significance was determined by ANOVA; p<0.01 = **, p<0.0001 = ****.

An essential characteristic of hRSV infection at the subcellular level is the formation of IBs that regulate the spatio-temporal compartmentalisation of viral RNA replication(29). smFISH imaging showed that hRSV-N and P are transcripts principally localised within cytoplasmic IB-like structures, as previously reported(27) **(Fig.5A).** Daprodustat treatment reduced the number of IBs, although treatment had no discernible impact on their average size **(Fig.5A)**. We assessed the intracellular levels of ISG15 in hRSV infected cells and noted significantly increased nuclear expression upon Daprodustat treatment in hRSV-N expressing cells **(Fig.5B).** These data support our earlier in vivo findings that Daprodustat enhances ISG15 and innate immune induction in hRSV infection, demonstrating the specificity of this finding in virus infected cells.

**Figure 5:**
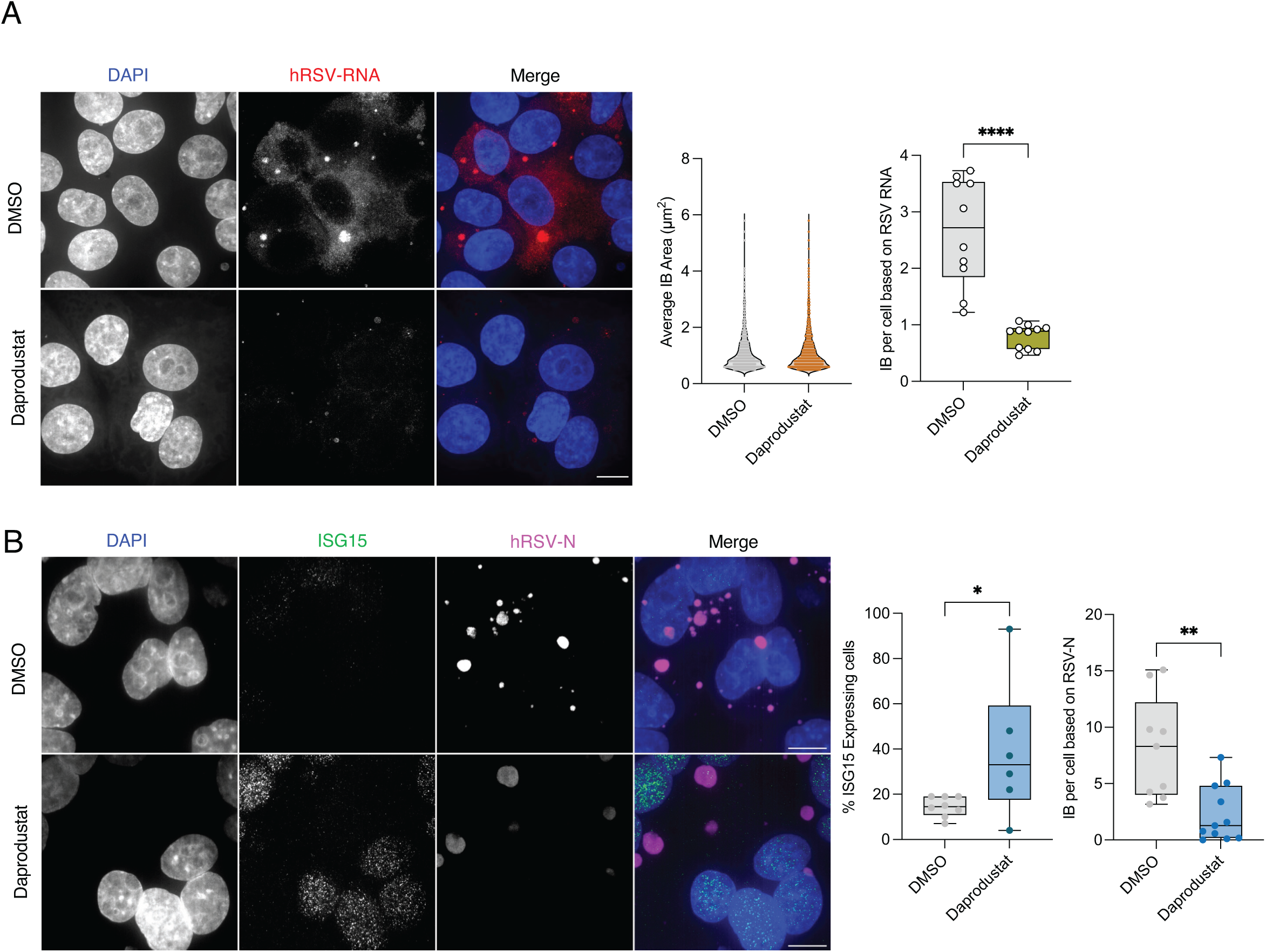
Daprodustat reduces the number of IBs and enhances nuclear ISG15 expression. **(A**) HEp-2 cells were infected with hRSV (MOI 1) and treated with Daprodustat (50µM) for 48h. Viral RNA was visualised through fluorescent *in situ* hybridisation using directly conjugated probes targeting hRSV-N and P transcripts. The area and number of viral IBs per cell was quantified. Individual channels are shown together with merged colour images, scale bar = 10µm. **(B)** ISG15 expression was visualised in hRSV infected HEp-2 cells treated with or without 50µM of Daprodustat by immunofluorescent staining. Cells were also stained for the hRSV-N protein to assess viral replication, scale bar = 10µm. The percentage of ISG15 positive cells per field of view was quantified, as was the average number of viral IB per cell. Statistical significance was determined by students t-test; p< 0.05 = *, p <0.01 = **.

### Daprodustat inhibition of hRSV is HIF dependent

To ascertain whether HIF directly impacts hRSV replication, we utilised the well characterised renal carcinoma cell line RCC4 which lacks functional VHL and constitutively expresses HIF-1α and 2α under normoxic conditions(65). Overexpression of VHL in the RCC4 cells (RCC4-VHL) restored physiological degradation of HIF-isoforms under normoxic conditions **(Fig.6A)**. GFP-hRSV replication was significantly reduced in RCC4 compared with RCC4-VHL cells, consistent with endogenous HIFs restricting viral replication **(Fig.6A-B)**. ISG15 gene and protein expression were increased in RCC4 cells following hRSV infection compared to RCC4-VHL cells **(Fig.6B, Supp Fig.6A)**. As an additional model, we used the independent renal cell line (786-0 cells), which lack both VHL and HIF-1α transcripts(65) allowing us to assess the role of HIF-2α to hRSV replication. We observed a similar reduction in hRSV N protein expression in the VHL expressing 786-0 cells **(Supp Fig.6B).** In line with our RCC4 results, ISG15 expression was increased following hRSV infection, although to a lesser degree. To extend these findings, we performed siRNA knockdown of both HIF-1α and 2α and assessed the effect of Daprodustat on viral replication in HIF depleted cells. Daprodustat had a reduced effect on viral replication when either HIF isoform was ablated **(Fig.6C),** confirming a role for both HIF-isoforms in regulating hRSV replication. Quantification of NDRG1 expression, a HIF-1α and 2α co-regulated gene(40), confirmed the functional ablation of HIF transcriptional activity **(Supp Fig.6C).** In summary, these data indicate that both HIF isoforms play direct roles in restricting hRSV replication.

**Figure 6:**
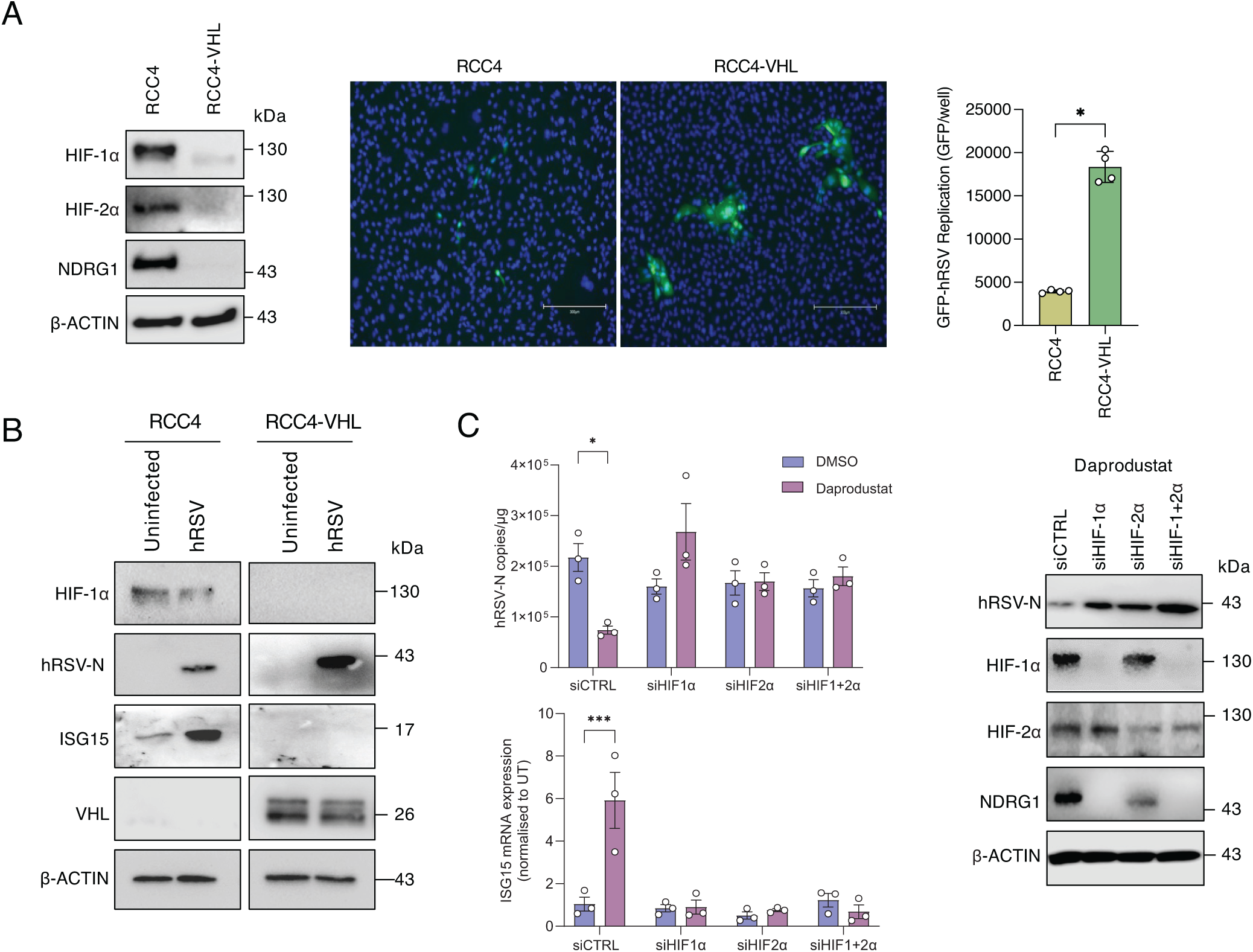
HIF-dependent restriction of hRSV. **(A)** RCC4 or RCC4-VHL cells were infected with GFP-hRSV at an MOI of 1 for 48h. Expression of HIF-1α, HIF-2α and NDRG1 was assessed by immunoblot and GFP expression quantified by Clariostar. **(B)** Expression of ISG15, hRSV-N, HIF-1α, VHL and β-actin was assessed by immunoblot. **(C)** Calu-3 cells were transfected with siRNA targeting HIF-1α or HIF-2α either individually or in combination followed by infection with hRSV and treatment with 50µM of Daprodustat. Viral replication was assessed 48h post-infection by RT-qPCR with quantification of hRSV-N transcripts and ISG15 mRNA. Protein expression of HIF-1α, HIF-2α NDRG1, hRSV-N and β-actin was determined by immunoblot. Unless otherwise stated, points represent independent biological replicates and are plotted as mean ± SD. Statistical significance was determined by ANOVA, p<0.01 = **, p<0.0001 = ****.

### Daprodustat exerts its antiviral effect on hRSV through RLR and type I IFN signalling

Given the augmentation of *ISG15* and other ISGs in our transcriptomic analysis, we assessed the impact of Daprodustat on IFN signalling in hRSV-infected Calu-3 cells over a 48h period. We noted increased expression of both ISG15 and IRF3 within 24h of hRSV infection in cells treated with Daprodustat, but limited modulation of these proteins in polyinosinic-polycytidylic acid (Poly-I:C) treated cells **(Supp Fig.6D)**. Using HEK 293-cells stably expressing a firefly luciferase reporter under the control of the IFN-β promoter (HEK-293-IFN-β-luciferase cells), we demonstrated that Daprodustat significantly increased reporter activity in hRSV-infected cells **(Supp Fig.6E)**. To assess a direct role for IFN-signalling in the antiviral activity of Daprodustat we treated cells with a competitive antagonist of the IFN-alpha receptor (IFNAR-Inh)(66). IFNAR-Inh treatment inhibited STAT-1 phosphorylation in response to IFNα treatment confirming efficacy in Calu-3 cells **(Supp Fig.7A)**. Combining the IFN-antagonist with Daprodustat limited its antiviral activity on hRSV N-gene expression and infectivity along with *ISG15 expression*, but with negligible changes in HIF mediated transcriptional activity **(Fig.7A, Supp Fig.7B)**. Quantification of the infectious viral titre confirmed the antiviral effect of Daprodustat in the presence of the INFAR-Inh (**Fig.7A)**. To extend these findings, we showed that hRSV infection of IFN-alpha receptor knock out cells (IFNAR KO) were insensitive to the antiviral activity of Daprodustat (**Supp Fig.7C**). When assessing the IFN-β promoter activity derived from the supernatants of these cells, we noted no Daprodustat-mediated increase in activity in hRSV-infected IFNAR-KO cells (**Supp Fig.7C**).

**Figure 7:**
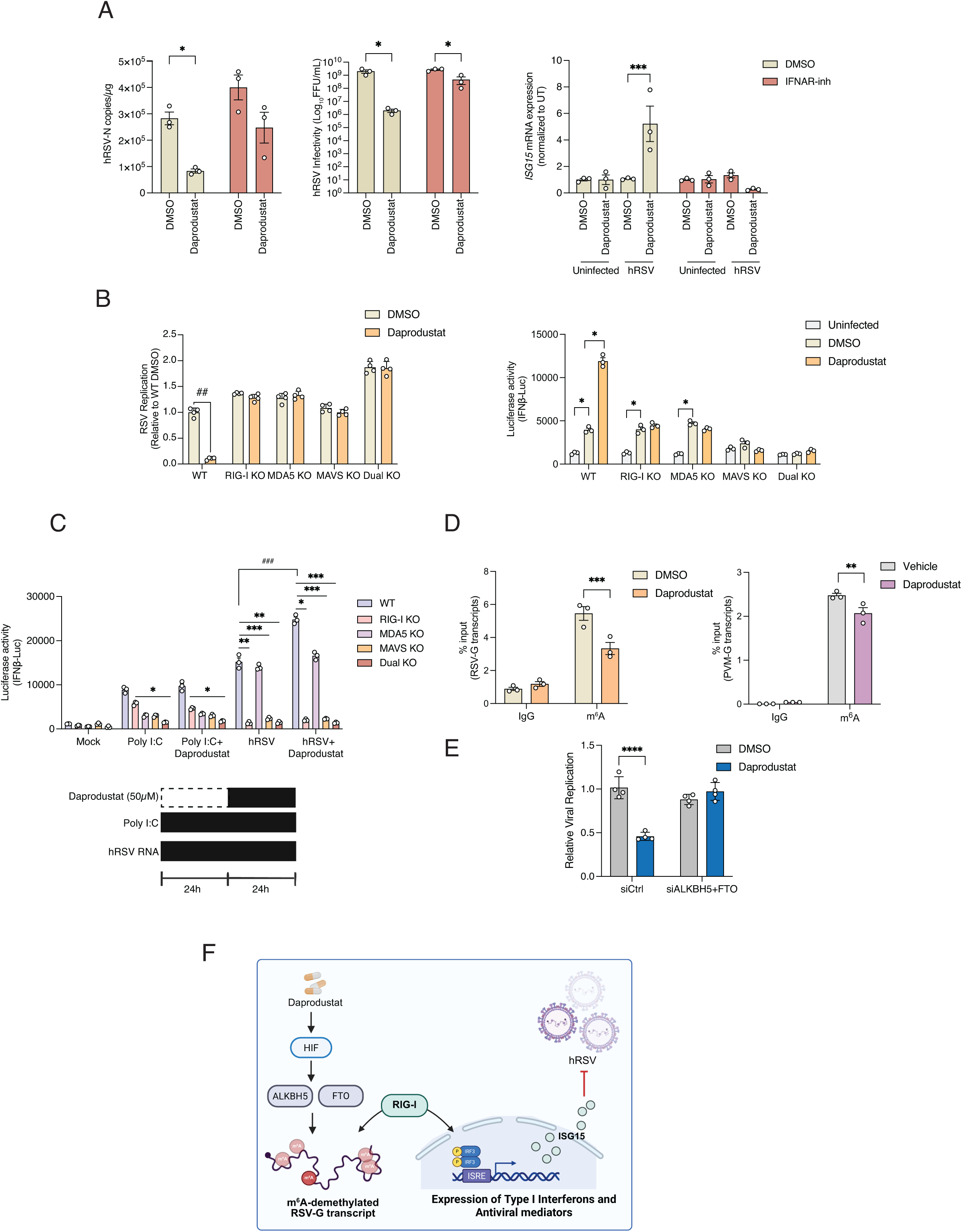
Daprodustat inhibition of hRSV replication is dependent on RIG-I signalling. **(A)** Calu-3 cells were infected with hRSV (MOI 1), treated with Daprodustat (50µM) and the IFN-alpha receptor inhibitor (10µM) for 48h. The infectious titre was assessed by FFU assay and hRSV-N and ISG15 mRNA expression was determined by RT-qPCR. **(B)** HEK293-IFN-β luciferase reporter cells were infected with hRSV (MOI 1) and treated with 50µM of Daprodustat. IFN-β-promoter luciferase activity and hRSV-N expression by immunofluorescence were quantified at 96h post-infection in WT and KO cells. **(C)** Quantification of hRSV and PVM-G transcripts from m^6^A RNA immunoprecipitation from hRSV infected Calu-3 or PVM infected lung tissue respectively, expressed as % input of total RNA. RNA input was normalised so an equal number of viral copies was used in the pull-down. **(D)** WT and KO cells were transfected with hRSV-particle associated viral RNA or poly I:C and treated with or without Daprodustat. Luciferase activity was quantified 48h post transfection. Data is representative of 3 biological replicates with statistical significance determined by ANOVA; p <0.05 = *, <0.01 = **, <0.001 = *** and <0.0001 = ****. **(E)** Calu-3 cells were co-transfected with siRNA against ALKBH5 and FTO, followed by infection with hRSV-GFP (MOI 1) followed by treatment with 50µM Daprodustat. Viral replication was assessed at 48h through quantification of the GFP by Clariostar. **(H)** Schematic showing the proposed mechanism through which Daprodustat modulates viral RNA methylation to promote increased innate sensing. **(F)** Schematic showing the proposed model of Daprodustat regulation of m^6^A methylation increasing innate sensing of hRSV.

As Daprodustat showed a reduced inhibition of hRSV infection in Vero cells **(Fig.4D)**, which do not produce endogenous IFN, we hypothesised that the antiviral phenotype occurs via cell-intrinsic PRRs such as RIG-I or MDA5. To assess this we infected HEK-293-IFN-β-luciferase WT, RIG-I, MDA5, double (RIG-I and MDA5) or MAVS KO cells(67)with a fluorescent GFP-hRSV, enabling the simultaneous measurement of viral replication and IFN induction. Increased levels of hRSV replication were observed in RIG-I and RIG-I/MDA5 dual KO cells compared to WT, demonstrating these receptors restrict viral replication, and while Daprodustat reduced viral replication in WT cells, this effect was not seen in cells lacking RIG-I, MDA5, or MAVS (**Fig.7B**). Quantification of IFN-β-driven luciferase confirmed increased activity in WT, RIG-I and MDA5 KO cells on hRSV infection, which was not observed in MAVS or RIG-I/MDA5 dual KO cells (**Fig.7B**). These data suggest that RIG-I and MDA5 play redundant roles in hRSV-induced activation of the IFN-β promoter in this setting. Notably, hRSV-infected WT cells treated with Daprodustat exhibited a significant augmentation in reporter activity that was absent in the KO cells, suggesting that RIG-I and MDA5 signalling play a key role in HIF regulation of innate viral RNA sensing pathways. To evaluate whether Daprodustat increased the activity of other PRRs such as cGAS-STING or TLR3 we treated our IFN-β reporter line with the cGAS-STING agonist diABZI or cGAMP and showed a dose-dependent activation that was not influenced by Daprodustat (**Supp Fig.7D**). Next, we considered if this was a generic anti-viral response and used Encephalomyocarditis virus (EMCV), a well-known activator of RLR signalling(68). Infection of our panel of KO cells with EMCV resulted in RIG-I and MDA5 dependent inductions of IFN-β promoter activity, which was not further enhanced by Daprodustat treatment (**Supp Fig.7E**). To examine whether Daprodustat increased detection of hRSV RNA through innate RNA sensing, we transfected our KO cells with RNA isolated from hRSV particles or with poly I:C, a positive control for RLR activation. hRSV-RNA activated the reporter in the WT cells and this effect was diminished in the RIG-I, MAVS and RIG-I/MDA5 dual KO cells but not in MDA5 KO cells, demonstrating an essential role for RIG-I to sense hRSV-RNA **(Fig.7C**). Daprodustat treatment of hRSV-RNA transfected cells significantly increased IFN-β reporter activity in both the WT and MDA5 KO cells, but not in cell lines ablated for RIG-I or MAVS expression **(Fig.7C).** Importantly, in poly I:C treated cells Daprodustat had minimal additional impact on the activation of the IFN-β reporter. Collectively, these findings support a role for Daprodustat to enhance the detection of hRSV RNA in a RIG-I dependent manner. Given the recent findings that loss of N6-methyladenosine (m^6^A) in hRSV-G transcripts resulted in increased detection of viral RNA by nucleic acid sensing(69) as well as HIF-regulated expression of the demethylase ALKBH5 reduces global m^6^A methylation(70), we hypothesised that HIF-driven depletion of RNA methylation would promote innate immune recognition of hRSV RNA. To examine this, we performed an m^6^A-RNA immunoprecipitation of RNA isolated from hRSV-infected Calu-3 cells treated with or without Daprodustat, noting a significant reduction in viral N gene expression post-infection (**Supp Fig.7F**). We observed an enrichment of m^6^A modified hRSV-G RNAs compared to NS1 and N transcripts, with Daprodustat treatment significantly reducing the levels of m^6^A methylated hRSV-G transcripts (**Fig.7D** and **Supp Fig.7G**). Further, we confirmed that the modulation of m^6^A methylation by Daprodustat extended to PVM, as significantly less m^6^A-modified PVM-G transcripts were precipitated from the total lung RNA extracted from vehicle and treated animals **(Fig.7D**). Given these data, we assessed whether Daprodustat retained antiviral activity against hRSV in the absence of the two main m^6^A demethylases, ALKBH5 and FTO(71, 72). Using siRNA knockdown of ALKBH5 and FTO in hRSV infected Calu-3 cells we ablated the antiviral activity of Daprodustat **(Fig.7E, Supp Fig.7H)**. Taken together, these findings demonstrate that the antiviral effect of Daprodustat is mediated through modulation of viral RNA m^6^A modification, promoting detection through the innate nucleic acid sensing pathway **(Fig.7F)**.

## DISCUSSION

This study demonstrates that activation of HIF-signalling by the PHDi Daprodustat limits pneumoviral replication *in vivo* and *in vitro,* through enhanced activation of innate sensing. Daprodustat treatment of PVM infected mice led to a significant reduction in pulmonary viral antigen expression, coupled with a reduced inflammatory response in the lungs. There is accumulating evidence that HIFs enhance intrinsic pathways to limit damage in acute lung injury through promoting blood vessel repair and the regeneration of alveolar type II pneumocytes(73, 74). Coupled with recent findings showing that HIFs play a key role in the differentiation of type I and II alveolar cells during embryonic development(75), implicate hypoxic signalling as a key protective pathway in the lung. Recent findings have demonstrated a protective influence of Vadadustat treatment, a new generation prolyl hydroxylase inhibitor, improving clinical outcomes in patients with severe lung injury resulting from SARS-CoV-2 infection as well as enhancing airway repair in murine models of infection(31). Analysis of the murine lung transcriptome revealed that Daprodustat promoted expression of numerous innate immune response genes in PVM infection, including interferon-stimulated genes such as *Isg15*. The increased *Isg15* expression in lung and spleen tissues further supports the immunomodulatory effects of Daprodustat and extends our earlier findings showing that HIF restricted nucleolin expression(44), demonstrating further roles for HIF to limit pneumovirus replication through modulation of innate sensing. Additionally, DCQ analysis of our transcriptomic data sets revealed modulation of specific immune cell sub-sets in the lung of Daprodustat treated animals infected with PVM. To improve the accuracy of these findings, application of either single-cell or spatial transcriptomic techniques would result in improved definition of the specific changes in the immune cell compartments induced by HIF signalling in pneumoviral infection. Using the natural mouse pathogen PVM to model pneumoviral disease allowed us to study viral replication in an *in vivo* setting that emulates many of the clinical features of severe hRSV infection, such as acute pneumonia, and leukocyte recruitment(57, 58). We observed that activation of HIF signalling limited both pulmonary viral antigen expression and immune cell recruitment compared with vehicle control groups. In addition, levels of *Isg15* mRNA in the spleen were significantly increased in PVM infected mice treated with Daprodustat. Protein modification by ISG15 (ISGylation) represents a significant element of the IFN-induced immune response, regulating critical antiviral factors important for the control of several viral infections such as Influenza and Sindbis virus(76, 77). Elevated expression of *ISG15* was previously reported in hRSV-infected nasopharyngeal washes and *in vitro* model systems(14), consistent with our observation in the lung and spleen of PVM-infected mice. ISG15 has been implicated as a post-translational regulator of HIF-1α expression, with ISGylation suppressing HIF-mediated gene transactivation and destabilising HIF in a proteasome-dependent manner(78). Notably, HIF-1α has been shown to induce transcriptional expression of *ISG15* through the presence of multiple hypoxic response elements in the *ISG15* promoter region(78, 79). Our transcriptomic analysis of infected PVM lungs revealed that HIF increased the expression of numerous innate immune response genes. Collectively these data suggest that in the context of pneumoviral infection, HIF acts in tandem with innate sensing to limit viral replication. This potential synergism may explain the marked increase in *Isg15* RNA expression in both the spleen and lungs of PVM infected animals and is supported by our confocal imaging data showing nuclear ISG15 staining in Daprodustat treated hRSV infected cells (**Fig.5B**). Importantly, using smFISH labelling of viral RNA combined with immunofluorescent imaging of the viral N protein, Daprodustat treatment limited the number of hRSV replication sites or IBs. We have previously demonstrated using this approach that activation of HIF limits the establishment of SARS-CoV-2 replication sites in early infection(42, 80). Whilst there are significant differences in the biology and replication cycles of coronaviruses and pneumoviruses, the restriction by HIF in their respective replication complex establishment may highlight common pathways that these viruses subvert to attain productive infection. These potential shared mechanisms could enhance our knowledge of virus-host interactions and contribute to more effective prevention and treatment strategies for respiratory viral infections.

In the lung HIF-1α is ubiquitously expressed in all pulmonary cell types(81), whilst HIF-2α is more commonly expressed in cells of the vascular endothelium and type II pneumocytes(82). Our siRNA and HIF-overexpression models allowed us to demonstrate a role for both HIF-alpha subunits to restrict viral replication indicating redundancy between either isoform. Blocking IFN signalling abrogated the antiviral activity of Daprodustat, indicating that the type I IFN pathway is required to limit viral replication, in agreement with the enhanced innate immune gene expression seen *in vivo*. Using ISG15 as a marker for innate gene activation, we found that ablation of HIF expression in the context of viral infection led to a reduction in ISG15 protein expression. In addition, disruption of RLR signalling significantly reduced the impact of Daprodustat on both viral replication and innate immune activation. RLRs are principal PRR that detects hRSV RNA in the early stages of infection to initiate the innate antiviral response. Previous studies have shown that genetic silencing of RIG-I *in vitro* had little impact on hRSV growth in lung epithelial cell lines(83), in agreement with our findings that showed a modest increase in replication in RIG-I knock-out cells. However, children with loss of function mutations in the *IFIH1* gene, which encodes MDA5, show higher susceptibility to hRSV-associated bronchiolitis, due to an inability to produce IFN-β(84), highlighting the importance of RLR sensing in the context of clinical infection. We observed that activation of the IFN-β promoter in response to hRSV infection was dependent on both RIG-I and MDA5, as only cells with ablated expression of both factors or of MAVS failed to mount a response to hRSV infection, indicating redundance between RIG-I and MDA5 (**Fig.7B**). Importantly, we demonstrated that enhancement of IFN-β promoter activity by Daprodustat in hRSV infected cells required functionality of both RIG-I and MDA5.

To dissect the mechanism by which HIF activation enhances innate immune sensing, we focused on the direct interaction between viral RNA and host PRRs. We found that viral particle associated RNA was sensed primarily by RIG-I when transfected into cells. Daprodustat treatment of viral-RNA, but not poly I:C, transfected cells resulted in increased IFN-β promoter activation, suggesting that HIF specifically promotes the detection of hRSV RNA rather than being a generic activator of RIG-I/MDA5 sensing. Importantly, neither RIG-I or MDA5 appear to be directly regulated by HIF and lack a hypoxic response element in their promoter regions. hRSV, like many RNA viruses, contains RNA modifications within its genome, the most common of which is N^6^-methyladenosine (m^6^A)(85) and there is an accumulating body of evidence suggesting that viruses acquire such modifications as a strategy to avoid host innate immunity(85, 86). In hRSV, m^6^A sites are concentrated in the G gene and depletion of these motifs results in increased immune recognition of viral RNA by RIG-I(69). Our findings highlight a previously undefined mechanism for HIF signalling to increase detection of pneumoviral RNA through modulation of RNA methylation. Whilst this work did not define the specific mechanism it reveals a potentially new area for exploration to leveraging RNA modification to promote innate immune signalling in pneumoviral infection.

In conclusion, this study identified Daprodustat as a promising host-directed antiviral agent against pneumoviruses, acting through HIF-dependent enhancement of cellular nucleic acid sensing. These findings not only suggest a potential new therapeutic approach for hRSV and related viral infections but also provide important insights into the molecular interplay between hypoxia signalling and antiviral immunity.

## METHODS

### Cells and Reagents

All cells, including co-cultures, were cultured at 37°C and 5% CO_2_ in a standard culture incubator and exposed to hypoxia using an atmosphere-regulated workstation set to 37°C, 5% CO_2_ and 1% O_2_ (Invivo i200, Baker-Ruskinn Technologies). Calu-3 cells were cultured in Advanced DMEM (Sigma-Aldrich) supplemented with 10% fetal bovine serum (FBS), 2mM L-glutamine, 100 U/mL penicillin and 10 mg/mL streptomycin (Invitrogen). HEp-2, Vero, RCC4, RCC4-VHL, 786-0, 786-0-VHL, BHK-21, HEK293-ISRE-Luc reporter, HEK293-P125 (IFNβ-Luc promoter cell) WT, RIG-I, MDA5, RIG-I/MDA5 KO and MAVS KO cells were cultured in DMEM (Sigma-Aldrich) with the same supplements as described above (see **Supp Table 1**). Human PBECs were purchased from Lifeline Cell Technologies. Airway epithelial cells were cultured in Airway Epithelial Cell medium (PromoCell, Heidelberg, Germany) in submerged culture. PBECs were cultured on PureCol-coated 0.4 μm pore polyester membrane permeable inserts (Corning) in serum-free airway epithelial cell media, brought to air-liquid interface, replacing basal media with Air-Liquid Interface Epithelial Differentiation Medium (Lifeline Cell Technologies). The media was exchanged every 2 days and apical surfaces washed with PBS weekly to disperse accumulated mucus, for a minimum of 6 weeks.

### Animals, PVM infection and Daprodustat treatment

Animal work was approved by the local University of Liverpool Animal Welfare and Ethical Review Body and performed under UK Home Office Project Licence PP4715265. Female BALB/c mice (BALB/cAnNCrl), 6-8 weeks of age were purchased from Charles River and maintained under SPF barrier conditions in individually ventilated cages. Mice were divided into two groups for treatment with vehicle or Daprodustat (n = 4 per group). Animals were treated with 30mg/kg of Daprodustat (MedChem Express) by oral gavage 2h pre-infection and then twice a day until sacrifice. Drug was dissolved in 99% double distilled H_2_O, 0.5% methyl cellulose and 0.5% Tween-80. Mice were intranasally infected with the PVM J3666 strain at a dose of 10^4 PFU under light ketamine anaesthesia. The PVM J3666 strain, (a kind gift of Prof Andrew Easton, University of Warwick) was passaged twice in mice. Lungs were harvested, and the supernatant titrated before in-vivo infection experiment Animals were euthanised by cervical dislocation on day 5 post-infection and tissues collected for, RNA viral titre and histological examination.

### Histological and immunohistological examination and RNA-ISH

The left lungs and spleen were fixed in 10% buffered formalin for 48 h, and stored in 70% ethanol until processing. Lungs (longitudinal section) and spleens (cross sections) were trimmed and embedded in paraffin wax. Consecutive sections (3 µm) were prepared and stained with haematoxylin and eosin (HE) for histological assessment and subjected to immunohistology (IH) for the detection of PVM-G antigen. IH was performed using the horseradish peroxidase (HRP) method. Briefly, after deparaffination, sections underwent antigen retrieval in citrate buffer (pH 6) for 20 min at 98°C, followed by incubation with mouse anti-PVM-G diluted in dilution buffer (Agilent Dako) overnight at 4°C. After blocking of endogenous peroxidase (peroxidase block, Agilent Dako) for 10 min at room temperature (RT), incubation) samples were incubated with rabbit anti-mouse IgG (Abcam) for 1 h at RT, followed by rabbit-HRP (Envision+System) for 30 min at RT in an autostainer (Agilent Dako). Sections were counterstained with haematoxylin. Further sections were stained for Iba1 (monocytes/macrophage marker) and CD3 (T cell marker) as previously described(87). To detect *ISG15* mRNA expression, RNA-ISH was performed using the RNAscope ISH method (Advanced Cell Diagnostics (ACD), Newark, California), Mus musculus ISG15 ubiquitin-like modifier (Isg15) (Mm-Isg15-O1; cross-detects Gm9706) oligoprobes, and the automated RNAscope 2.5 Detection Reagent Kit (brown) according to the manufacturer’s protocol, and as previously described(88). To determine whether monocytes/macrophages and neutrophils express *ISG15* mRNA, a combined RNA-ISH/IHC protocol was applied after established immunohistology protocols for Iba and Ly6G (neutrophil marker) proved to maintain immunolabeling after the ISH pretreatment steps (baking, deparaffinizing, incubating in RNAscope Hydrogen Peroxide, cooking in RNAscope 1X Target Retrieval Reagents solution, and incubating in RNAscope Protease Plus). Sections were then subject to the RNA-ISH protocol, followed by staining for Iba1 and Ly6G, respectively, combining previously published protocols(87, 88).

### RNA-seq analysis

Right upper lung lobes from PVM infected mice at 3dpi were homogenised in 1 ml of Trizol reagent (Thermofisher) using a QT tissue lyser and stainless-steel beads (Qiagen) at 50 oscillations for 5 minutes. The homogenates were clarified by centrifugation at 12,000xg for 5 min before full RNA extraction was carried out according to manufacturer’s instructions. RNA was quantified and quality assessed using a Nanodrop (Thermofisher) before a total of 1µg was DNase treated using the TURBO DNA-free™ Kit (Thermofisher) as per manufacturer’s instructions. mRNA expression of ISG15 and IFN-b measured using 1step QuantiNova® SYBR® Green RT-PCR Kit (Qiagen) and normalised against mouse 18S ribosomal RNA reference gene. RNA was sequenced using a 300bp paired-end Illumina sequencing protocol (Novogene, UK). Host reads were mapped to the human transcriptome and differential expression analysis performed using DESeq2. Pathway analyses were performed using the enrichGO package in Bioconductor using R-Studio version 2024.12.0. Relative cell quantities were estimated using Digital Cell Quantification (DCQ)(61) implemented in the immunedeconv R package (v2.1.0). Briefly, gene counts were normalised using the DESeq2 package (v1.42.1) and input into immunedeconv with the function deconvolute_mouse (normalised_counts, “dcq”). Statistical differences across experimental groups were assessed using a Kruskal–Wallis test followed by Dunn’s post hoc test for pairwise comparisons, implemented via the rstatix package (v0.7.2). Results were visualised using ggplot2 (v3.5.2). Data is available through the following accession code GSE290159.

### hRSV and PVM in vitro propagation

Human hRSV subtype A (A2 strain) was grown in HEp-2 cells and cell lysates concentrated using Vivaspin® 20, 10,000 MWCO PES columns (Sartorius) as reported previously(44). Briefly, cells were infected at MOI 0.2 for 2h, the inoculum removed, cells washed with PBS and cultured in DMEM containing 5% FBS. After 5 days, cells were harvested and cellular lysates clarified and concentrated using a Vivaspin® 20 column. GFP-hRSV was propagated using the same method as described for hRSV(44). BHK-21 cells were cultured in DMEM containing 10% FBS and infected with PVM (J3666 strain) at MOI 0.0025 at 33 °C in 5% CO_2._ After 10 days, the cell lysates concentrated using 10,000 MWCO PES columns (Sartorius). The hRSV, GFP-hRSV and PVM viral stocks were aliquoted, snap frozen in liquid nitrogen and stored at-80°C.

### Plaque assays

To quantify infectious hRSV, samples from infected cells and lung homogenates from infected mice were serially diluted 1:10 and used to inoculate monolayers of HEp-2 cells for 2h. Inocula were then replaced with DMEM containing 1% FCS with a semi-solid overlay of 1.5% carboxymethyl cellulose (Sigma-Aldrich). Cells were incubated for 4-5 days, fixed in 4% PFA, stained with 0.2% crystal violet (w/v) and plaques enumerated, where the limit of detection of this assay was 10 focus forming units PFU/mL. PVM quantification in BHK-21 cells was conducted using a modified version of the standard immunofluorescent plaque assay described by Watkiss et al. (2013) (58). Lung samples from in-vivo infections were serially diluted tenfold in DMEM medium, incubated for 72 h at 33°C with 5% CO_2_, and subsequently fixed in ice cold methanol and acetone and stained with PVM-G monoclonal antibody at room temperature. Plaques were visualized and counted using a fluorescent microscope (Zeiss).

### Immunoblotting

Cell lysates were prepared by washing cells with phosphate buffered saline (PBS), then lysing in RIPA lysis buffer (20 mM Tris, pH 7.5, 2mM EDTA,150 mM NaCl, 1% NP40, 0.1% SDS and 1% sodium deoxycholate) supplemented with Complete TM protease inhibitor cocktail (Roche) at 4°C for 5 min, followed by clarification by centrifugation (3 min, 12,000 rpm). Supernatants were mixed with Laemmli sample buffer, and boiled at 100°C, separated by SDS-PAGE and proteins transferred to polyvinylidene difluoride membrane (Immobilon-P, Millipore). Membranes were blocked in 5% milk in PBS/ 0.1% Tween-20, incubated with anti-HIF-1α (BD Transduction Laboratories, clone#610959), anti-NDRG1 (Cell signalling, clone#5196) or anti-β-actin (Sigma, clone#A5441) primary antibodies and appropriate HRP-conjugated secondary Mouse (DAKO, clone#P0447) or Rabbit antibodies (Cytiva, clone#NA934V). Chemiluminescence substrate (West Dura, 34076, Thermo Fisher Scientific) was used to visualize proteins using a ChemiDoc XRS+ imaging system (BioRad). Densitometric analysis was performed using FIJI (NIH).

### Confocal immunofluorescence microscopy and smFISH

Infected cells were fixed with 4% paraformaldehyde (PFA; Sigma) in PBS for 15 min, blocked with 20mM glycine in PBS and permeabilized with 0.5% Triton X-100 in PBS for 5 min. Samples were incubated with anti-hRSV-F primary antibody for 1h at room temperature, washed and incubated with Alexa Fluor secondary antibodies (Life Technologies) for a further hour at room temperature. After washing, slides were mounted with Fluoromount G (SouthernBiotech) containing 4′,6-diamidino-2-phenylindole (DAPI) for nuclei staining. smFISH was carried out as previously reported(80). Briefly, HEp-2 cells grown on #1.5 round-glass coverslips in a 24-well plate were fixed in 4% paraformaldehyde (ThermoFisher) for 30min at room temperature. Cells were permeabilised in PBS/0.1%

Triton X-100 for 10min at room temperature, followed by washes in PBS and 2X SSC. Cells were pre-hybridised in pre-warmed (37°C) wash solution (2×SSC, 10% formamide) twice for 20min at 37°C. smFISH probes designed against the hRSV-N and P transcripts were diluted to 500nM in hybridisation solution (2×SSC, 10% formamide,10% dextran sulphate) and incubated overnight at 37°C. Coverslips were then washed for 20min in pre-warmed wash solution at 37°C followed by counterstaining with DAPI (1μg/mL) diluted in wash solution. Cells were washed once with wash solution for 20min at 37°C and twice with 2XSSC for 10min at room temperature. Cells were imaged on an Olympus SoRA spinning disc confocal microscope. Quantification was performed using ImageJ software (Fiji). The average intensity of hRSV-RNA smFISH signal was defined based on an automatic threshold with a Minimum Error algorithm. The number of cells was quantified based on the DAPI signal. A watershed step was performed to separate nuclei in close proximity.

### siRNA silencing

Scramble, HIF-1α and HIF-2α siRNA were transfected to Calu-3 cells separately or in combination using DharmaFect 4 (Thermo Fisher). 24 h after transfection, Calu-3 cells were treated with DMSO (50µM) or Daprodustat (50 µM) for 48 h after inoculation with hRSV (MOI 1) and at 48hpi cells collected for RT-qPCR.

### ISRE and IFN-β Luciferase assay

HEK293-ISRE-Luc reporter cells were infected with hRSV at an MOI of 1 for 2h, the viral inoculum was removed and the cells cultured with growth media including Daprodustat (50µM). 24h post-infection, luciferase expression was assessed using a Firefly luciferase assay system (Promega, E1500) according to the manufacturer’s instructions. HEK293-p125 WT, RIG-I, MDA5, MAVS, RIG-I/MDA5 KO cell lines were seeded into 96 well plates and transfected with 150ng of viral RNA or poly I:C (Merck, P1530, Polyinosinic–polycytidylic acid sodium salt) complexed with Fugene 6 (Promega) per well. 24h after transfection, Daprodustat (50µM) was added to each well for 48h. Luciferase activity was determined as described above. For extraction of hRSV particle associated RNA, hRSV stocks were treated with RNAse I (Thermo) to remove any non-particle associated RNA. RNase was inactivated by heating at 95°C for 10mins and RNA extracted using TRIzol (Ambion) according to manufacturer’s instructions. Post-extraction RNA was quantified using a Nanodrop 2000 spectrophotometer (Thermo Scientific) and 150ng transfected into HEK293-P125 cells as described above.

### RT-qPCR

RNA was extracted from infected cells using the RNeasy Mini kit (QIAGEN), according to the manufacturer’s instructions. 500 ng of RNA were used to generate cDNA using an UltraScript cDNA synthesis kit (PCR Biosystems) and mRNA expression measured using the SyGreen Blue, SYBR green qPCR kit (PCR Biosystems). For quantification ΔC_t_ values were defined as the difference between the target gene Ct value and the Ct value of a housekeeper gene.

### RNA m^6^A immunoprecipitation

Total RNA was extracted from hRSV (MOI 1) infected Calu-3 cells treated with or without Daprodustat using the RNeasy Mini Kit (QIAGEN). The RNA was subsequently treated with TURBO DNase (Thermo Fisher Scientific) and further purified using the RNeasy Mini Kit. RNA concentration was measured, and equal copy numbers of hRSV RNA were incubated with Protein G Agarose beads (Cell Signaling) pre-bound and either Mouse IgG (SantaCruz) or anti-m^6^A Monoclonal antibody (Proteintech) in MeRIP buffer (50 mM Tris-HCl (pH 8.0), 150 mM NaCl, 0.1% NP-40, 1 mM EDTA) supplemented with RNase inhibitor (Promega). m^6^A-modified RNA was eluted using 6.7 mM m^6^A sodium salt, purified using Qiagen RNA extraction kit and performed RT-qPCR analysis.

### Quantification and statistical analysis

All data are presented as mean ± SD. P values were determined using a Mann-Whitney test or students t-test (two group comparisons) or a Kruskal–Wallis ANOVA (multi group comparisons) using PRISM version 10. In the figures * denotes p < 0.05, ** < 0.01, *** <0.001 and **** <0.0001.

## Acknowledgements

JH, ST and PACW are funded by the Chinese Academy of Medical Sciences Oxford Institute, CAMS Innovation Fund for Medical Sciences (CIFMS) funding code:2024-I2M-2-001-1. The McKeating laboratory is funded by a Wellcome Investigator Award 200838/Z/16/Z, Wellcome Discovery Award 225198/Z/22/Z, Chinese Academy of Medical Sciences Innovation Fund for Medical Science, China (grant number: 2018-I2M-2-002). The Stewart laboratory is supported by UK Medical Research Council (MRC) grants MR/W021641/1, MR/W005611/1, Innovate UK grant TS/W022648/1, Biotechnology and Biological Sciences Research Council grant BB/W020351/1 and Wellcome Trust grant 223733/Z/21/ZST. STa is supported by funding from the Biotechnology and Biological Sciences Research Council (UKRI-BBSRC) [grant number BB/T008784/1]. DM is funded by UKRI grants MR/W005611/1/ and MR/Y004205/1. J.R. acknowledges funding from the UK Medical Research Council [MR/Y013212/1; J.R.]. The authors are grateful to the technical staff in the Histology Laboratory, Institute of Veterinary Pathology, Vetsuisse Faculty, University of Liverpool, for excellent technical support.

## Supplementary Figure Legends

**Supplementary Figure 1:**
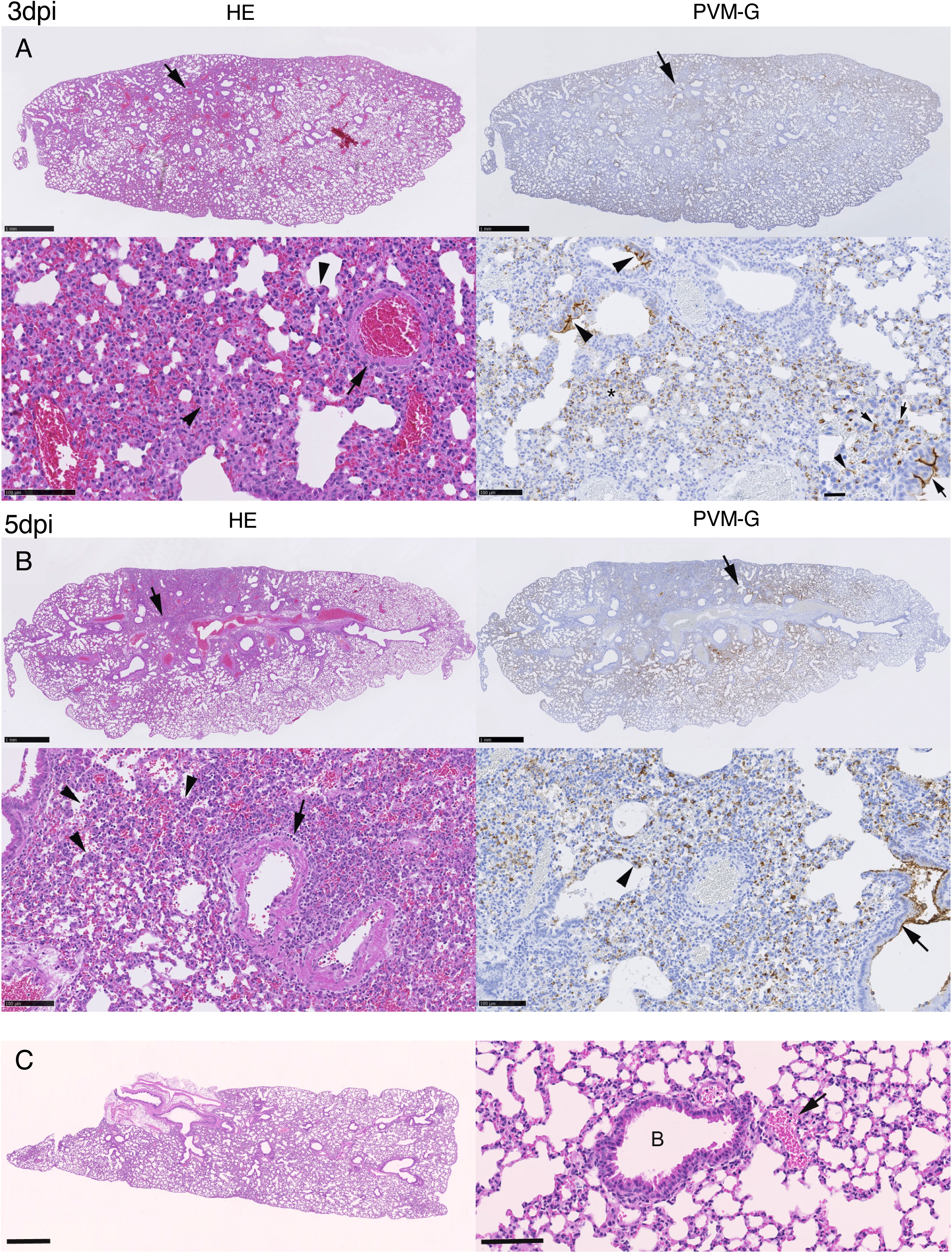
Pulmonary histopathology of PVM infected mice. Histological features and viral antigen expression were assessed at 3 and 5 dpi. **(A)** At 3 dpi, the lung parenchyma showed focal areas of increased cellularity (overview: arrow). Arrowheads of the higher magnification image show mild increases in interstitial cellularity and large activated type II pneumocytes in HE stained sections. Consecutive sections were stained for PVM-G that showed widespread expression (right image - overview). Large arrowheads on the bottom inset image indicate abundant infected type I pneumocytes, small arrowheads show infected type II pneumocytes in an area of increased cellularity (asterisk). The inset image bottom right, shows PVM-G expression in patches of respiratory epithelial cells, mainly along the luminal cell borders (arrowheads)**. (B)** At 5 dpi, focal dense areas of the parenchyma are indicated by the arrow on the overview images (HE stains). Arrowheads on the bottom left higher magnification image indicate the presence of desquamed alveolar macrophages/type II pneumocytes and some leukocytes in alveolar lumina and mild leukocyte recruitment and perivascular accumulation (arrow). Viral infection is widespread, with abundant viral antigen expression in alveoli (type I and II pneumocytes; arrowhead) and in respiratory epithelial cells in a few bronchioles. The abundant unstained cells are infiltrating leukocytes. The arrow in the overview image highlights the area depicted in the higher magnification. C) Lung, mock infected mouse. The parenchyma is unaltered, as shown in the overview (left) and in a closer view of a bronchiole (B) with adjacent vessel (arrow), surrounded by unaltered alveoli. Representative images are shown, and scale bars represent 50µm.

**Supplementary Figure 2:**
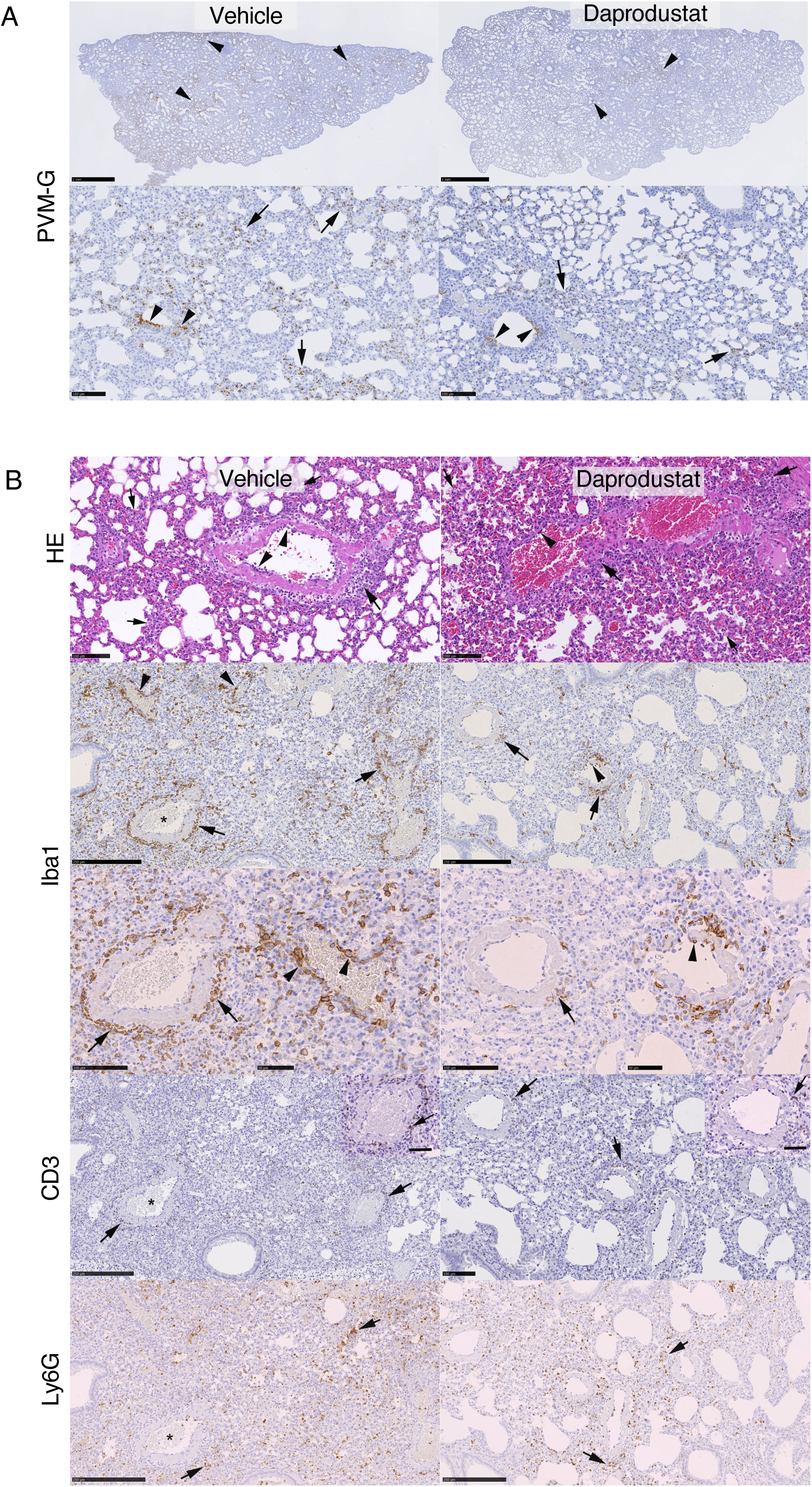
Histological features of Daprodustat treated PVM infected mice. **(A**) Immunohistology of viral antigen (PVM-G) in longitudinal sections of the left lobe of the lung at 3dpi. Arrowheads indicate areas of widespread viral antigen expression in the overview images (top row). Arrowheads in the higher magnification images (bottom row) highlight viral antigen expression in bronchiolar respiratory epithelial cells and arrows indicate expression in large groups of alveoli with abundant infected type I and II pneumocytes. **(B)** Immunohistology of longitudinal sections of the lung at 5 dpi of a vehicle treated (left column) and Daprodustat treated animal (right column). Arrowheads in the HE stained sections (top layer) identify muscular veins with leukocytes rolling along the endothelial cells, showing evidence of leukocyte recruitment into the tissue. Large arrows indicate examples of perivascular accumulation of mononuclear cells and small arrows alveoli with desquamed alveolar epithelial cells/alveolar macrophages in the lumen. Consecutive sections were stained for the monocyte/macrophage marker Iba1 (second and third layer). Arrows indicate areas of perivascular infiltrates and arrowheads show areas where monocytes/macrophage are emigrating from and accumulating around vessels. Macrophages are far less abundant in the lungs of Daprodustat treated mice where they are also present in the minimal perivascular infiltrates (arrows) accompanied by evidence of monocyte rolling (arrowhead). T cells (CD3+: bottom layer) are present in the lungs of both groups of mice in low numbers. The arrows point at T cells in perivascular infiltrates (insets: higher magnification). Neutrophils (Ly6G+; bottom layer) are present in the lungs of both groups of mice in low numbers. They are mainly seen in the focal areas of alveolar damage and inflammatory infiltration (arrows). Representative images are shown, and scale bars represent 50µm.

**Supplementary Figure 3:**
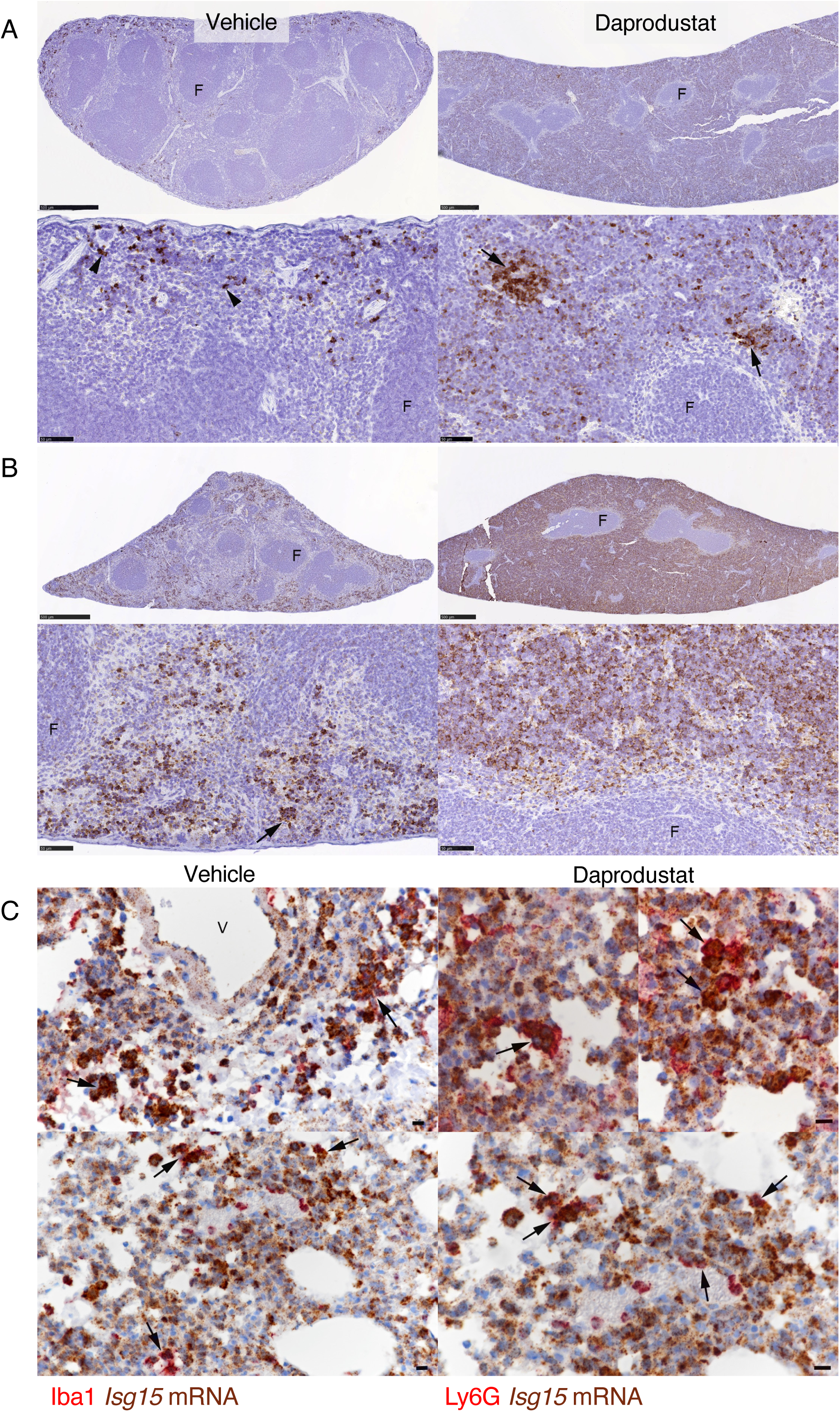
ISG15 gene expression in the spleen and infiltrating leukocytes in the lung after PVM infection. **(A)** Detection of *Isg15* mRNA expressing cells in the spleen of untreated (left column) and Daprodustat treated (right column) mock infected mice (top layers). In untreated mock infected mice, individual *Isg15* mRNA positive cells are present in the red pulp (arrowheads). In Daprodustat treated mock infected mice, where the red pulp is of much higher cellularity, numerous *Isg15* mRNA positive cells are observed. These form variably sized aggregates (arrows). **(B)** In untreated mice after PVM infection, the number of *Isg15* mRNA positive cells is substantially increased. Again, these positive cells form aggregates (arrow). RNA-ISH, with DAB as chromogen and haematoxylin counterstain. F: follicle. Bars = 25 µm in higher magnifications. **(C)** Lungs from untreated animals after PVM infection (5 dpi). Identification of *Isg15* transcripts in infiltrating leukocytes. Macrophages (Iba1+) with *Isg15* mRNA signal are abundant in a perivascular infiltrate (left: arrows). A higher magnification identifies *Isg15* mRNA in Iba1 positive alveolar macrophages and recruited, infiltrating macrophages (right: arrows). Neutrophils (Ly6G+) with *Isg15* mRNA signal are present in the parenchymal inflammatory infiltrates (left: arrows). The higher magnification highlights the nuclear morphology of the *Isg15* mRNA positive neutrophils, confirming that Ly6G co-expressing cells are neutrophils (right: arrows). *Isg15* RNA-ISH, with DAB as chromogen, followed by immunohistology with AEC as chromogen, haematoxylin counterstain V: vessel. Representative images are shown, and scale bars represent 10µm.

**Supplementary Figure 4:**
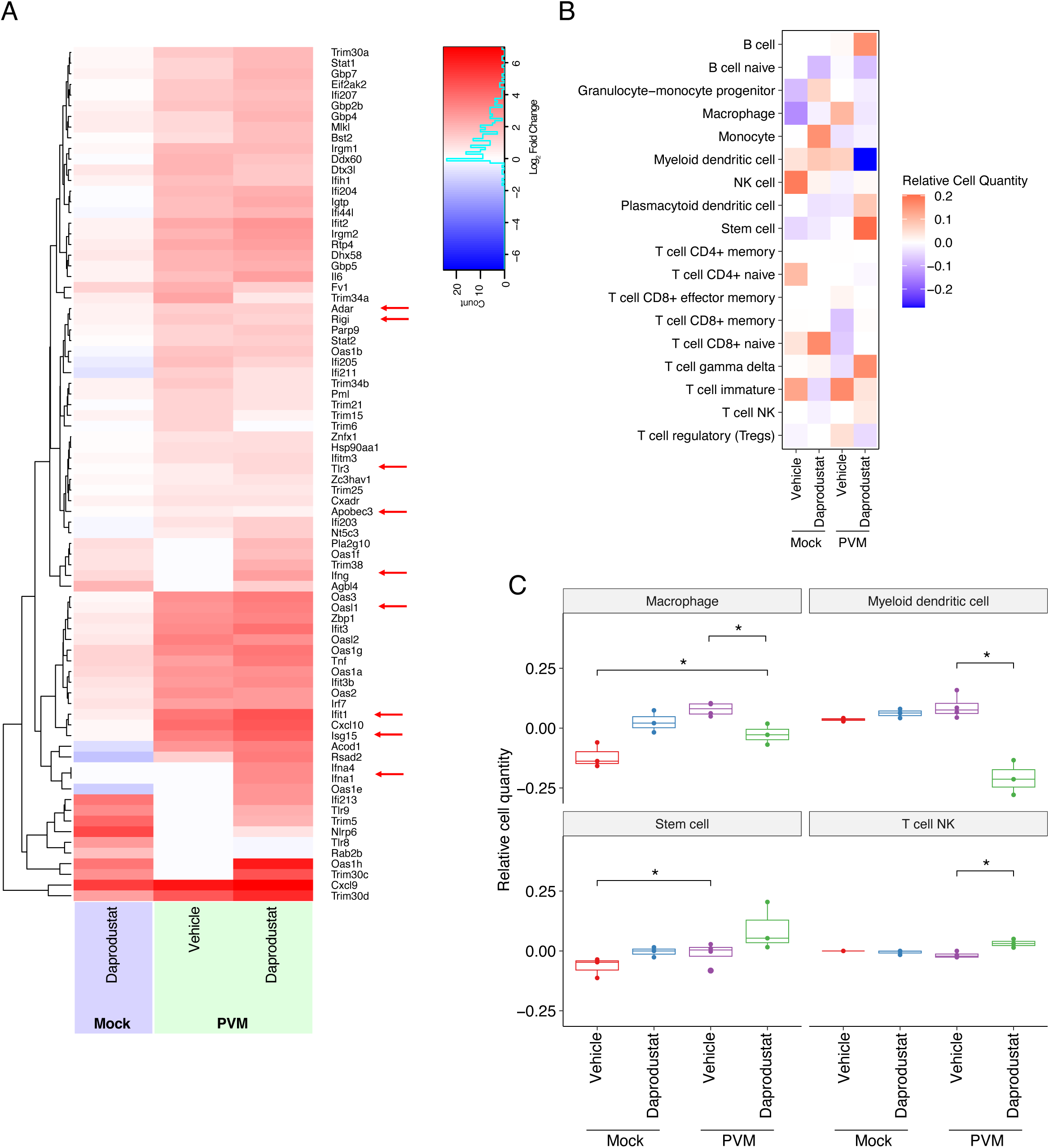
The impact of Daprodustat treatment on innate immune gene expression in uninfected and PVM infected mice. **(A)** Hierarchical clustering of innate response genes in murine pulmonary transcriptomes either treated with Daprodustat or infected with PVM at 3dpi and treated with either vehicle or Daprodustat compared to uninfected animals. Arrow denotes key innate immune genes and colours are derived from the log_2_ fold change in expression relative to uninfected, untreated animals. **(B)** Heatmap showing the average relative abundance of immune cell types estimated by Digital Cell Quantification (DCQ) from normalised gene counts using the immunedeconv package. Cell types are shown on the y-axis and experimental groups are on the x-axis. Values represent the mean relative cell quantity per group. **(C)** Boxplots showing the distribution of relative abundance for a given cell type across four experimental groups: uninfected and PVM infected mice treated with or without Daprodustat. Statistical differences were assessed using Kruskal–Wallis tests followed by Dunn’s post hoc tests with Benjamini– Hochberg correction. Asterisks indicate significant pairwise comparisons (*p < 0.05, **p < 0.01).

**Supplementary Figure 5:**
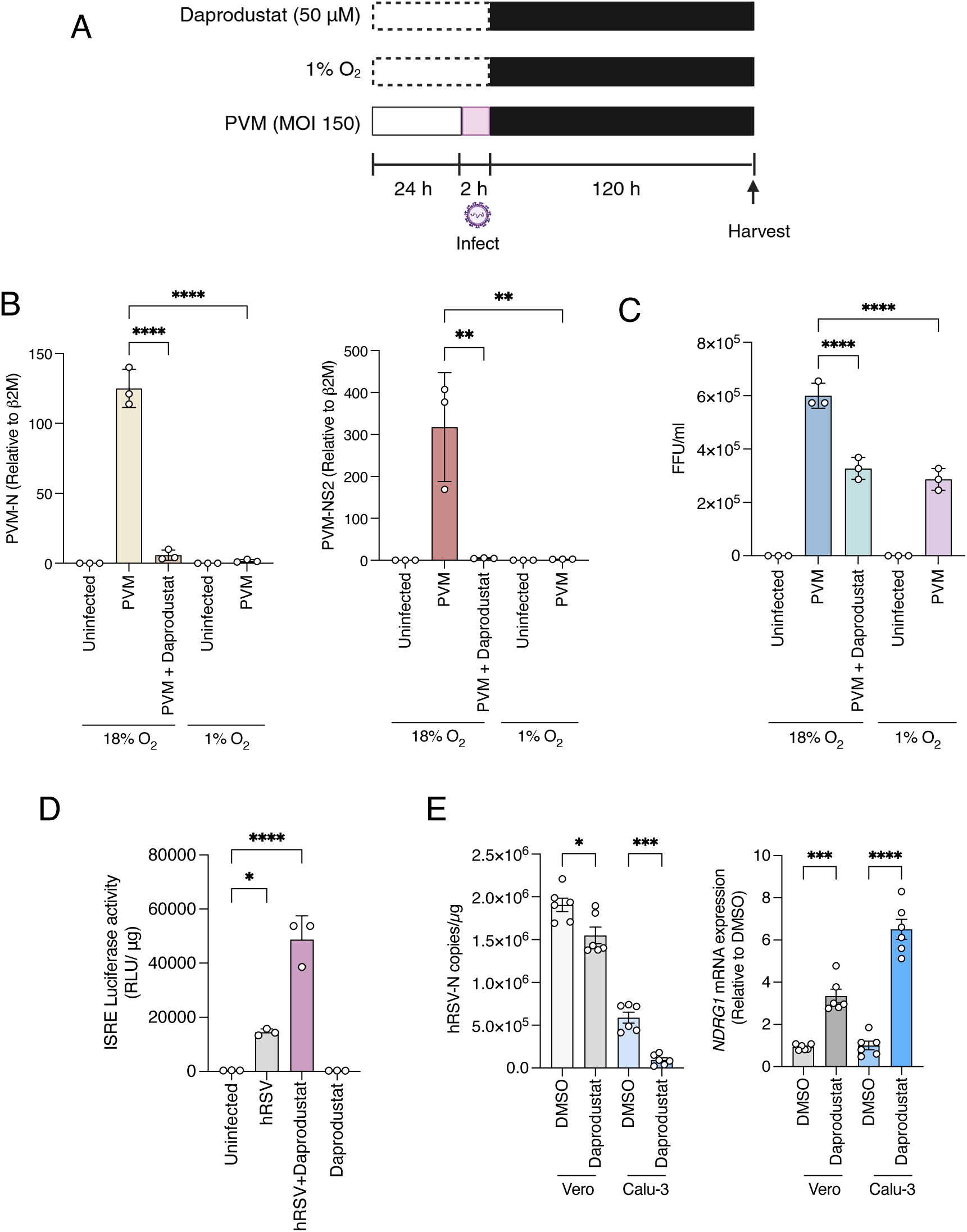
Daprodustat restricts PVM replication in vitro and induces innate sensing activity during hRSV infection. **(A)** Experimental schematic of PVM infection of BHK cells, where cells were infected at an MOI of 150 for 2h followed by removal of the inoculum. Cells were subsequently incubated in either 1% O_2_ or treated with 50µM of Daprodustat for 5 days. **(B)** Quantification of PVM N and NS2 transcripts by RT-qPCR, expressed relative to the uninfected control. Data is from three biological replicates and is plotted as mean ± SD. Statistical significance was determined by ANOVA; p<0.01 = **, p<0.0001 = ****. **(C)** At 5 dpi, PVM Infectious titre was assessed by FFU assay using an antibody against PVM-G. Data is representative of triplicate biological replicates and plotted as mean ± SD. Statistical significance was determined by ANOVA, p<0.01 = **, p<0.0001 = ****. **(D)** HEK293-ISRE-reporter cells were treated with supernatants from hRSV (MOI 1) infected HEp-2 cells with or without Daprodustat (50 µM). At 24 hpi, cell lysates were measured with Luciferase activity. **(E)** Vero or Calu-3 cells were infected with hRSV (MOI1) and treated with 50µM Daprodustat for 48h and hRSV-N or NDRG1 expression mRNA measured by RT-qPCR (mean ± SEM, n = 6, ANOVA, p<0.05 = *, p<0.001 = ***.)

**Supplementary Figure 6.**
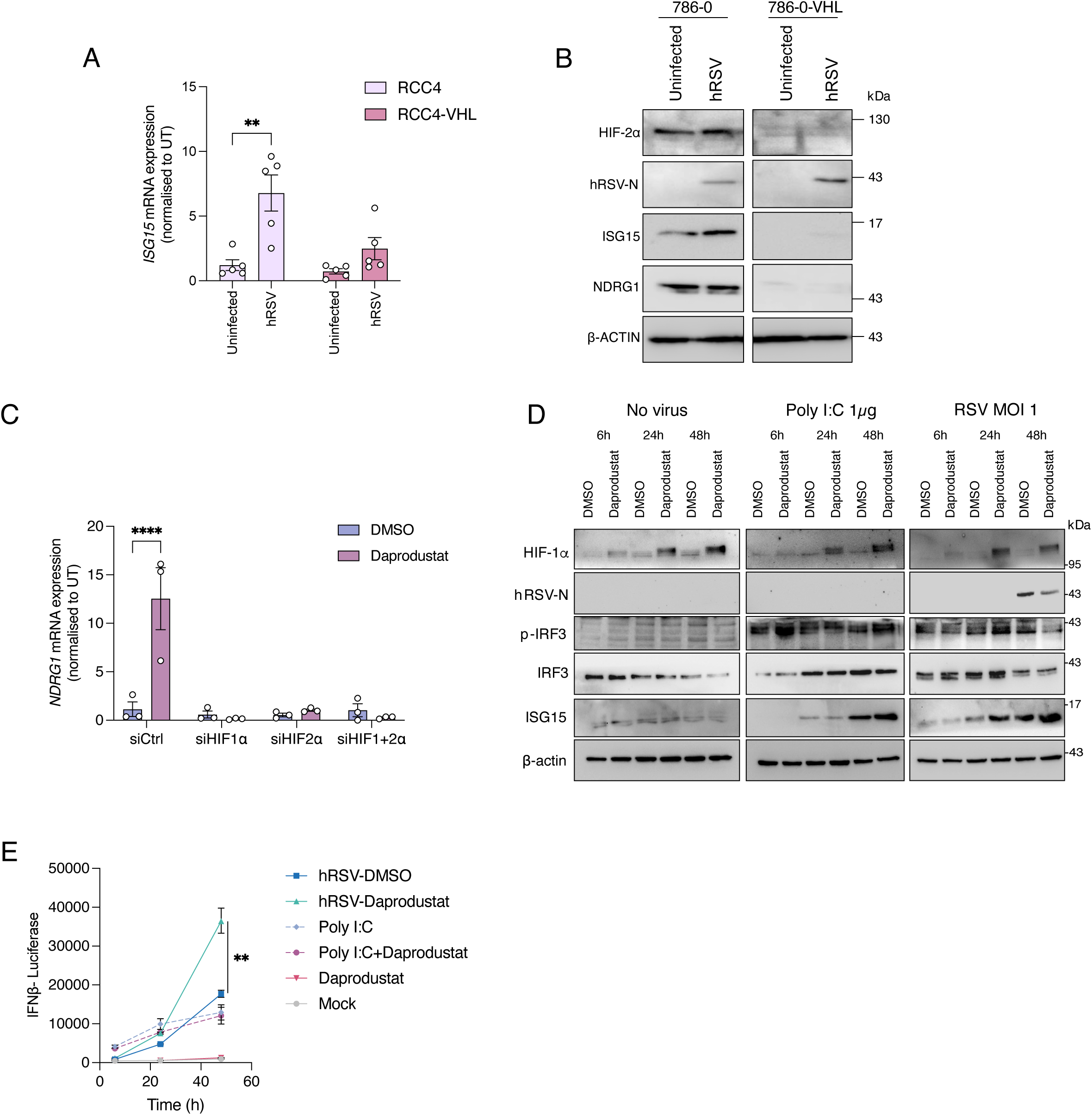
ISG15 expression is HIF-dependent. **(A**) *ISG15* gene expression in mock and hRSV infected RCC4 and RCC4-VHL cells. **(B)** 786-0 and 786-0-VHL cells were infected with hRSV for 48h (MOI 1) and cell lysates probed for hRSV-N, ISG15, HIF-2α, NDRG1 and β-actin expression by immunoblot. **(C)** *NDRG1* gene expression in Calu-3 cells transfected with siRNAs against HIF-1α and HIF-2α. **(D)** Expression of IRF3 and ISG15 was assessed in Calu-3 cells treated with stimulated with Poly-I:C or infected with hRSV (MOI 1) followed by treatment with Daprodustat 2h post infection or treatment. Cells were sampled at 6, 24 and 48h post treatment and expression of indicated proteins assessed by western blot. **(E)** HEK293-IFN-β luciferase reporter cells were infected with hRSV at an MOI of 1 followed by treatment with 50µM of Daprodustat. Luciferase reporter activity was quantified at the indicated timepoints over the course of 48h from both infected and uninfected cells treated with DMSO or Daprodustat.

**Supplementary Figure 7.**
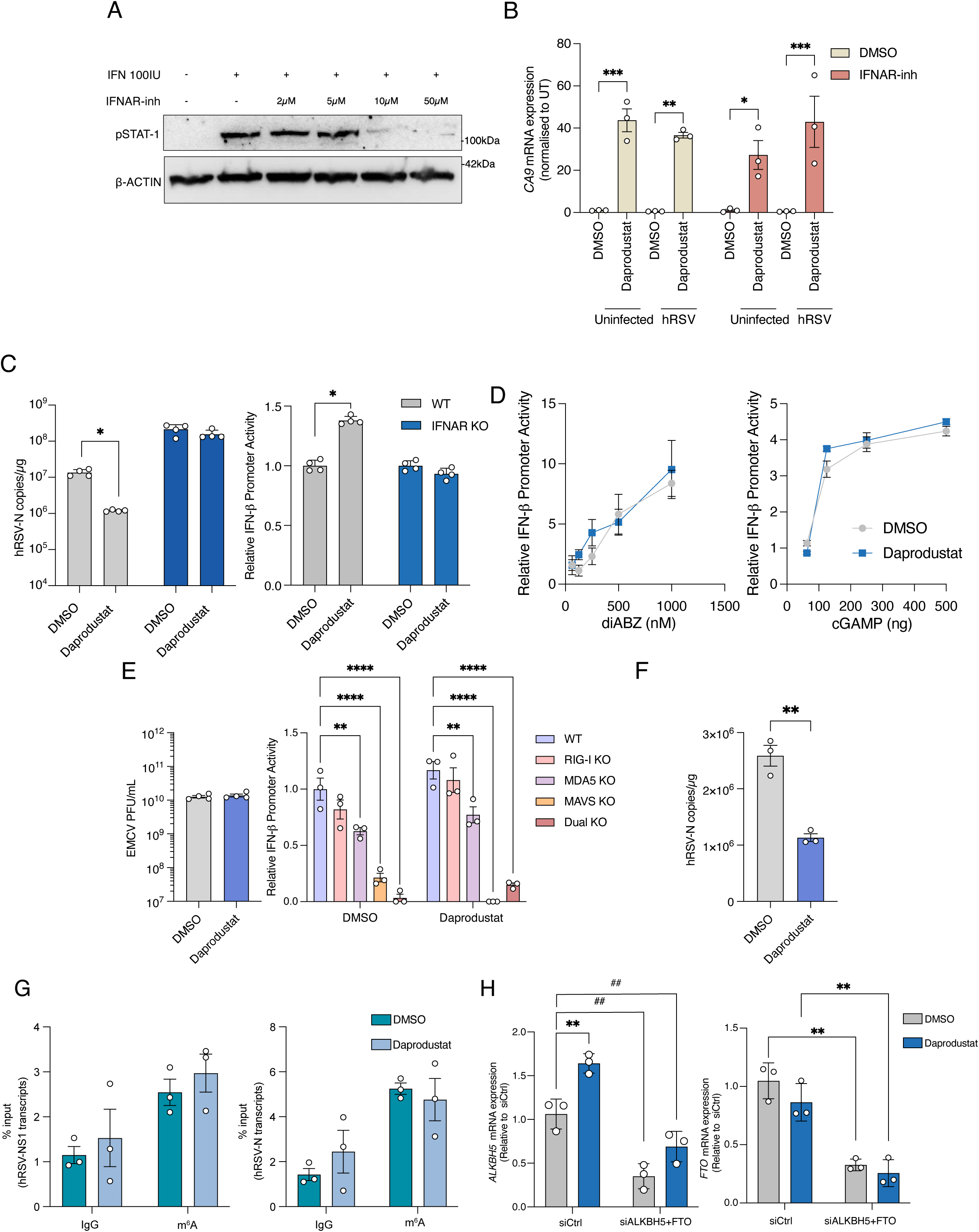
The antiviral effect of Daprodustat is mediated through modification of RNA m^6^A methylation. **(A)** Phosphorylated STAT-1 expression in Calu-3 cells treated with IFNα (100U) and increasing dose of IFNAR-inhibitor. **(B)** *CA9* gene expression in mock and hRSV infected Calu-3 cells treated with or without Daprodustat (50µM) plus the INFAR-inhibitor (10µM) for 48h. **(C)** HEK-293 WT or IFNAR knock out cells were infected with hRSV at an MOI of 1 followed by treatment with 50µM of Daprodustat for 48h. Viral replication was assessed through quantification of hRSV-N transcripts. Supernatants from this experiment were used to treat HEK293-IFN-β luciferase reporter cells with reporter activity expressed as relative to the DMSO control. **(D)** HEK293-IFN-β luciferase reporter cells were treated with increasing doses of diABZ or cGAMP with or without 50µM of Daprodustat. Luciferase activity was assessed 48h post-treatment and data expressed as relative to the DMSO control **(E)** Quantification of infectious EMCV from 293 cells treated with or without Daprodustat. Measurement of IFN-β luciferase from WT and indicated KO cells, infected with EMCV treated with or without 50µM of Daprodustat. Data is expressed as relative to WT cells treated with DMSO **(F)** qPCR quantification of hRSV copies from infected Calu-3 cells treated with or without Daprodustat. **(G)** Quantification of methylated hRSV N and NS1 RNA transcripts from m^6^A immunoprecipitation from total cellular RNA. **(H)** mRNA expression of *ALKBH5* and *FTO* in Calu-3 cells transfected with siRNAs targeting both genes.

---

**Supplementary Table 1:**
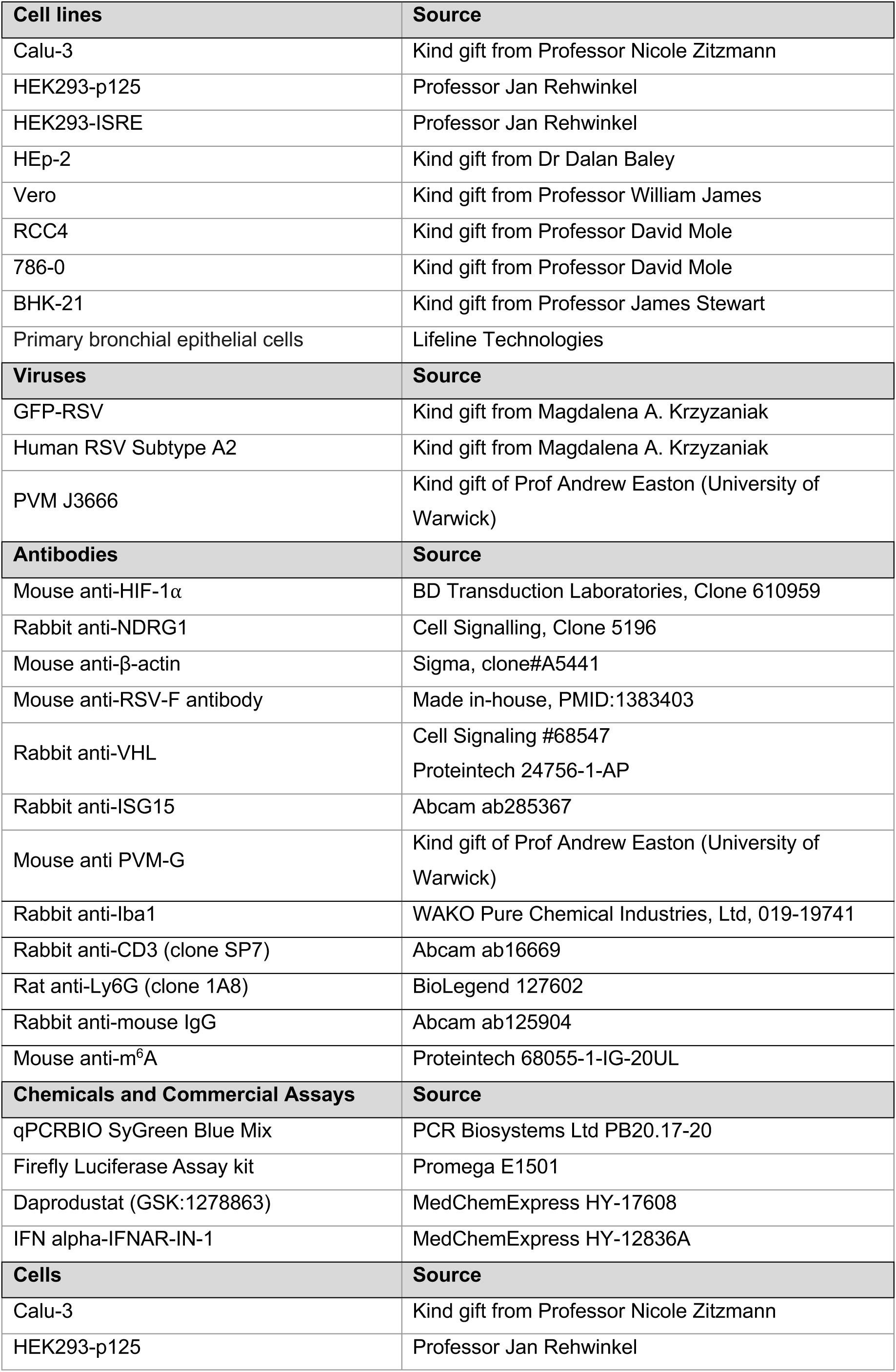
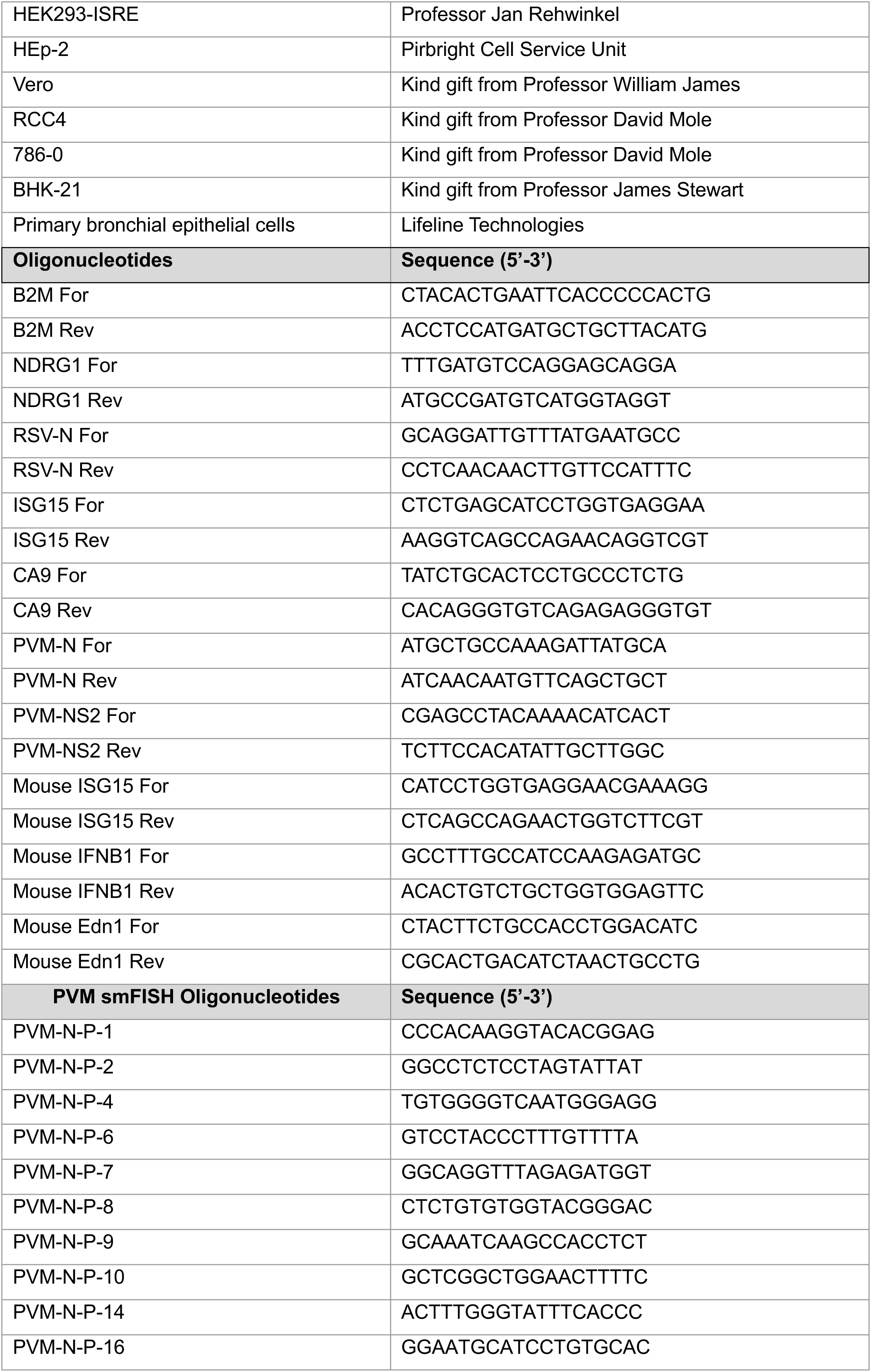
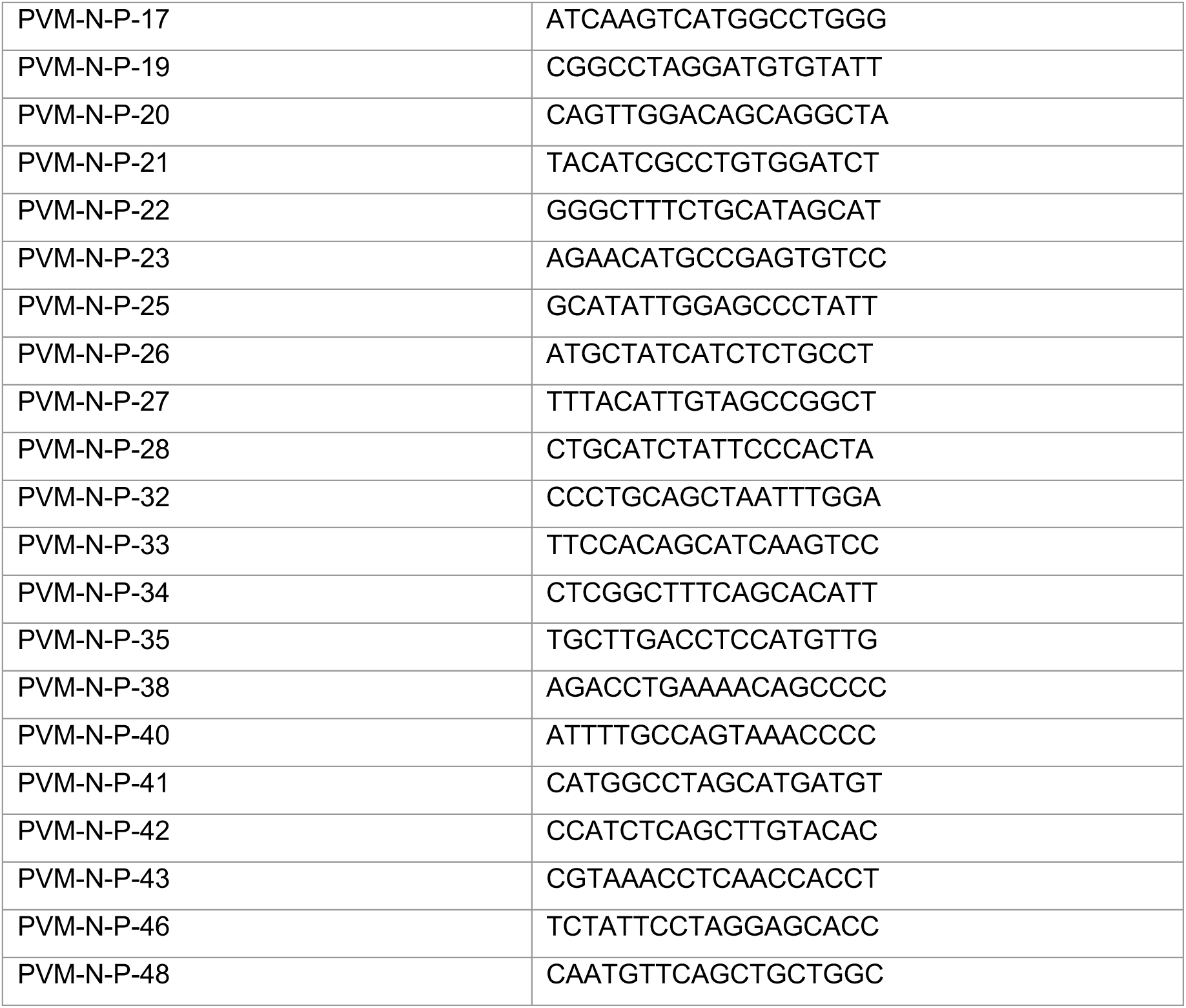
Key Resources.

